# Conditional frequency spectra as a tool for studying selection on complex traits in biobanks

**DOI:** 10.1101/2024.06.15.599126

**Authors:** Roshni A. Patel, Clemens L. Weiß, Huisheng Zhu, Hakhamanesh Mostafavi, Yuval B. Simons, Jeffrey P. Spence, Jonathan K. Pritchard

## Abstract

Natural selection on complex traits is difficult to study in part due to the ascertainment inherent to genome-wide association studies (GWAS). The power to detect a trait-associated variant in GWAS is a function of frequency and effect size — but for traits under selection, the effect size of a variant determines the strength of selection against it, constraining its frequency. To account for GWAS ascertainment, we propose studying the joint distribution of allele frequencies across populations, conditional on the frequencies in the GWAS cohort. Before considering these conditional frequency spectra, we first characterized the impact of selection and non-equilibrium demography on allele frequency dynamics forwards and backwards in time. We then used these results to understand conditional frequency spectra under realistic human demography. Finally, we investigated empirical conditional frequency spectra for GWAS variants associated with 106 complex traits, finding compelling evidence for either stabilizing or purifying selection. Our results provide insight into polygenic score portability and other properties of variants ascertained with GWAS, highlighting the utility of conditional frequency spectra.

## Introduction

Over the last two decades, genome-wide association studies (GWAS) have uncovered countless associations between genetic variants and complex traits. In the process, it has become apparent that the effect sizes of trait-associated variants are inversely correlated with their frequencies (Speed *et al*., 2012; Yang *et al*., 2010, 2015). This relationship has been interpreted as a sign that natural selection acts on complex traits (Schoech *et al*., 2019; Zeng *et al*., 2018, 2021). Furthermore, patterns of diversity around trait-associated variants appear to deviate from neutral expectations (Gazal *et al*., 2017; Speed *et al*., 2020).

To explain these observations, different models of selection on complex traits have been proposed. Several groups have proposed models of stabilizing selection, wherein intermediate trait values are favored (Lande, 1976; Simons *et al*., 2018; Turelli, 1984). Stabilizing selection on a trait results in selection against the minor allele at independent genetic variants underlying the trait (Robertson, 1956; Simons *et al*., 2018; Walsh and Lynch, 2018). In other words, the direction of selection at individual alleles depends on the frequency of the allele. In contrast, others have proposed a model of purifying selection, which entails selection against all new mutations (Caballero *et al*., 2015; Eyre-Walker, 2010; Keightley and Hill, 1990; Zeng *et al*., 2021). Under such models, the direction of selection is independent of allele frequency. Finally, some have proposed models of directional selection on complex traits, where the direction of selection at individual alleles depends on the sign of their effects (Guo *et al*., 2018).

Distinguishing between competing models of selection is essential for interpreting the results of GWAS. The mode and strength of selection acting on traits determines whether GWAS tend to ascertain variants with large or small effects. Moreover, because selection influences the genetic architecture of traits at the molecular level, selection can also shape the extent to which discovered variants are pleiotropic and further upstream in regulatory networks, or more specific and further downstream in regulatory networks (Mostafavi *et al*., 2023). Naturally, this has consequences for the proportion of heritability explained by discovered variants (O’Connor *et al*., 2019; Weiner *et al*., 2023), as well as the portability of polygenic scores, or the extent to which a polygenic score derived from a particular GWAS cohort is predictive in genetically dissimilar individuals (Durvasula and Lohmueller, 2021; Wang *et al*., 2020; Yair and Coop, 2022). Thus, a clear understanding of selection on complex traits is not only essential for studying complex trait evolution, but also for understanding complex trait architecture more broadly.

Selection on complex traits has been challenging to investigate in part because of the technical limitations of GWAS. First, the power to ascertain a trait-associated variant in GWAS depends on the variant’s frequency and effect size; GWAS are more powered to ascertain variants with large minor allele frequencies and large effect sizes. For traits under selection, the effect size of a variant determines the strength of selection on the variant, which in turn constrains its frequency. This logic also applies to traits that are genetically correlated with traits under selection. Trait-associated variants ascertained in GWAS are therefore enriched for common variants and thought to be depleted for the most strongly selected variants (Manolio *et al*., 2009; Pritchard, 2001). Second, GWAS generally rely on genotype arrays and imputation panels to infer genotypes genome-wide. By implicitly prioritizing a subset of genetic variation, arrays and imputation algorithms will bias discovered variants in a manner that is particularly difficult to model (Clark *et al*., 2005; Lachance and Tishkoff, 2013). As a result, the process of GWAS ascertainment can obscure the hallmark signatures of selection.

To avoid the biases induced by GWAS ascertainment, we propose conditioning on the allele frequency in the GWAS cohort by leveraging additional, diverged populations. These additional populations provide orthogonal sources of information about selection. Moreover, conditioning on the frequency in the GWAS cohort effectively conditions on GWAS ascertainment itself. Yet, in order to combine information across populations, we need to develop a framework for describing how selection and non-equilibrium demography impact the distribution of allele frequencies in one population conditional on another.

A wealth of previous work has used population genetics theory to answer questions at the intersection of selection, demography, and the (joint) allele frequency spectrum. Maruyama (1974) derived theoretical expectations for the impact of selection on allele frequencies within a single population under equilibrium demography. More recently, others have characterized the impact of non-equilibrium demographies on both neutral variation (Baharian and Gravel, 2018; Bhaskar and Song, 2014; Bhaskar *et al*., 2015; Myers *et al*., 2008; Ragsdale *et al*., 2018; Rosen *et al*., 2018; Schraiber, 2018; Spence *et al*., 2016; Terhorst and Song, 2015) and selected alleles within a single population (Chen and Slatkin, 2013; Evans *et al*., 2007; Schraiber *et al*., 2016; Slatkin, 2001; Song and Steinrücken, 2012; ΁ivković and Stephan, 2011). Consequently, we have a solid theoretical expectation for how selection and demography impact the allele frequency spectrum within one population. There has also been a line of work exploring how these two evolutionary forces impact the joint frequency spectrum for two or more populations (Dilber and Terhorst, 2024; Gutenkunst *et al*., 2009; Jouganous *et al*., 2017; Kamm *et al*., 2017, 2020; Kern and Hey, 2017; Lukić *et al*., 2011; Lukić and Hey, 2012; Yang *et al*., 2014).

However, the effect of conditioning on frequencies in one population has been under-explored. We propose conditioning on frequencies in a GWAS cohort as a means of studying selection on complex traits, but previous work on conditional frequency spectra has focused on neutral variants, not selected variants (Harpak *et al*., 2016; Durvasula and Sankararaman, 2020). While it is known that different models of selection leave distinct signatures on joint frequency spectra, it is unclear if this also holds for conditional frequency spectra. Conditional frequency spectra require considering selection both forwards and backwards in time: first backwards from the conditional population to a shared ancestor, then forwards to the second, unconditioned population. The impact of selection on allele frequency trajectories backwards in time is poorly understood, especially under non-equilibrium demographies.

Thus, we sought to characterize conditional frequency spectra across modern human populations under different models of selection. Through our theoretical analyses of conditional frequency spectra, we identified strong qualitative signatures associated with each model of selection. We then used 106 complex trait GWAS to investigate empirical conditional frequency spectra at trait-associated variants and assess evidence for different models of selection on complex traits. Finally, we used this framework to understand how selection and non-equilibrium demography impact the portability of polygenic scores. Our results highlight the utility of conditional frequency spectra as a tool for studying selection on complex traits.

## Results

### GWAS ascertainment biases allele frequency spectra

We first sought to demonstrate the impact of GWAS ascertainment on the allele frequency spectrum. We began by curating a set of variants associated with 106 complex traits and diseases. We obtained 18,229 trait-associated variants from published GWAS of 94 quantitative traits in the UK Biobank “White British” cohort, approximately 337,000 individuals (http://www.nealelab.is/uk-biobank). We additionally obtained a set of 1,040 trait-associated variants from published GWAS of 12 complex diseases in various cohorts with European ancestries (Table S1).

We compared the derived allele frequencies of trait-associated variants to those of approximately 11 million non-associated variants imputed in the UK Biobank. As expected, we found that the frequencies of trait-associated variants are enriched for common variation, reflecting the bias induced by the process of ascertainment (Figure 1A). The tremendous discrepancy between the frequencies of trait-associated variants and the true allele frequency spectrum emphasizes the utility of conditional frequency spectra (Figure 1B). Similar to joint frequency spectra, conditional frequency spectra can encode information about demography and natural selection. Unlike joint frequency spectra, conditional frequency spectra can circumvent the biases induced by GWAS ascertainment by conditioning on the frequencies of trait-associated variants in the GWAS cohort.

**Figure 1:**
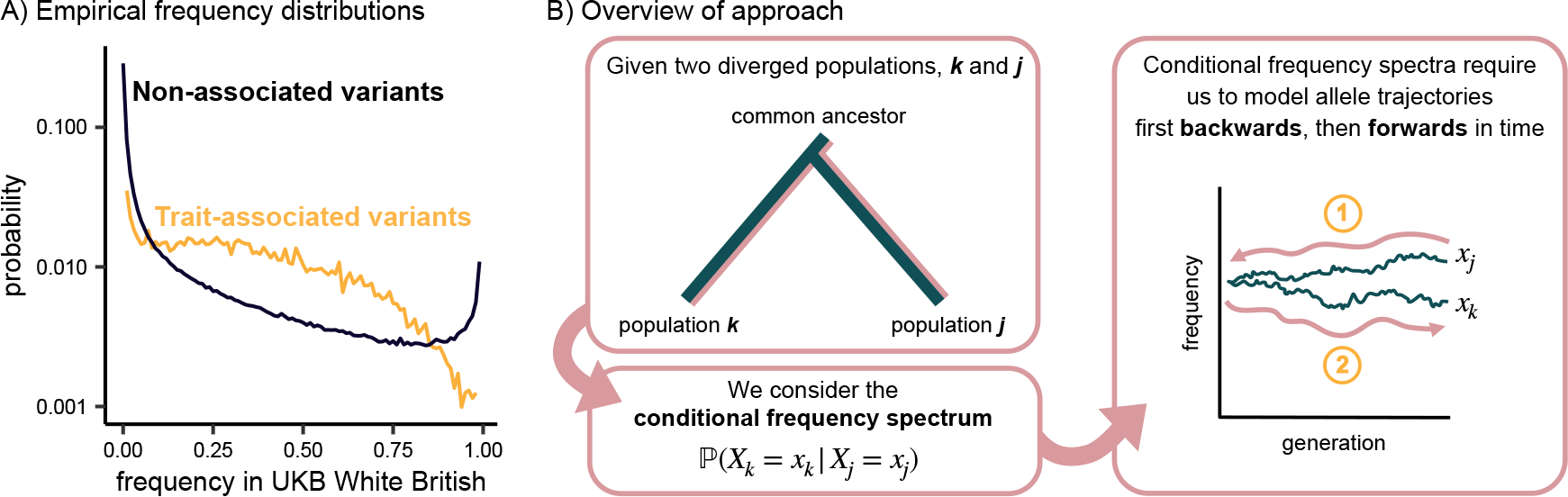
Overview. **A)** Derived allele frequency distribution of 19,269 trait-associated variants and approximately 11 million non-associated variants in the UK Biobank White British cohort. **B)** Overview of conditional frequency spectra. Given two populations *k* and *j*, the conditional frequency spectrum ℙ(*X_k_* = *x_k_* |*X_j_* = *x_j_*) requires us to first consider allele trajectories backwards in time from *x_j_* to the frequency in the common ancestor of populations *k* and *j*; then ultimately forwards in time to *x_k_*.

### Characterizing allele frequency dynamics over time

To build intuition for the allelic dynamics captured by conditional frequency spectra, we characterized the impact of demography and selection on allele frequencies over time. We considered four different demographic models relevant for human populations: equilibrium demography (i.e. constant population size), bottleneck, exponential population growth, and bottleneck followed by exponential growth. We also considered three types of selection: purifying selection against new alleles, directional selection on traits, and stabilizing selection on traits. These three modes of selection can be well-approximated by different allelic dynamics. Purifying selection acts as negative genic selection against derived alleles; directional selection at the trait level acts as positive genic selection for trait-improving alleles; and stabilizing selection can be approximated by perfect underdominance where only heterozygotes have reduced fitness (Robertson, 1956; Walsh and Lynch, 2018).

We first recapitulated well-known results about how selection impacts allele frequency trajectories forwards in time. We simulated allele frequencies under a Wright-Fisher model for 2,000 generations and a constant population size with *N_e_* = 10,000. Regardless of an allele’s frequency, positive selection and negative selection consistently act to increase and decrease frequencies, respectively. In contrast, allelic dynamics under stabilizing selection exhibit a frequency dependence because stabilizing selection results in selection against the *minor* allele. Alleles at low frequencies (e.g. 0.1) have nearly indistinguishable dynamics under stabilizing selection and negative selection (Figure 2A), but as the derived allele frequency increases, the dynamics under stabilizing selection gradually diverge from those under negative selection. When alleles are at a frequency of 0.5, dynamics under stabilizing selection resemble those of neutrality (Figure 2B), and at frequencies well above 0.5, stabilizing selection begins to look similar to positive selection (Figure 2C).

**Figure 2:**
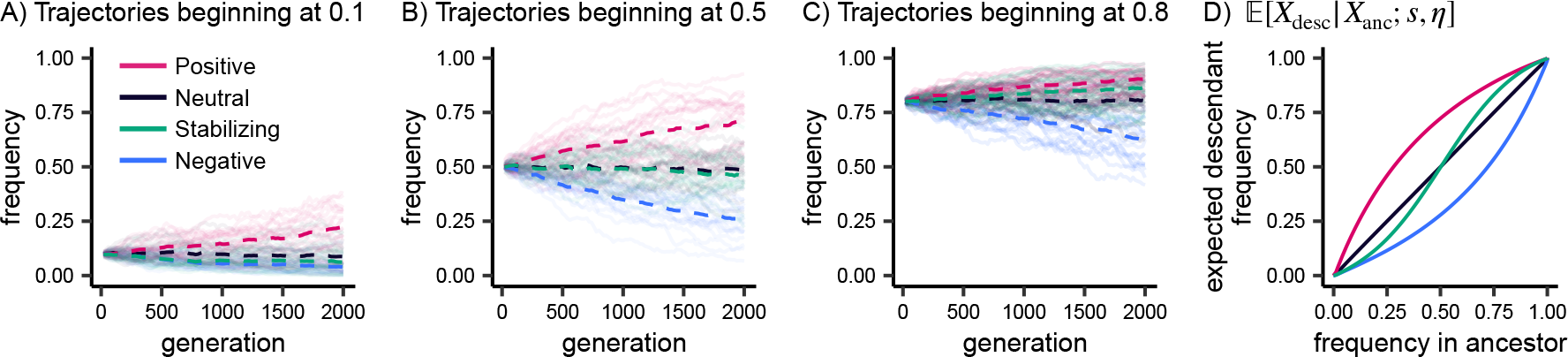
Forward transitions. **A, B, C)** Allele frequency trajectories simulated under a Wright-Fisher model. Dashed lines depict the mean frequency across 20 trajectories for each model of selection. **D)** Expected frequency in a descendant population conditional on the frequency in the ancestral population, computed with fastDTWF. For all panels, the demographic model consists of a constant population size with *N_e_* = 10,000 for 2,000 generations, and selection coefficients correspond to |*hs*| = 5.0 × 10*^−^*^4^.

We next summarized allelic dynamics by computing the probability of an allele transitioning from a given ancestral frequency, *x*_anc_, to a given descendant frequency, *x*_desc_, *P* (*X*_desc_ = *x*_desc_|*X*_anc_ = *x*_anc_; *s, η*), where *s* represents the selection scenario and *η* the demography. We computed these forward-in-time transition probabilities using two different approaches: Wright-Fisher simulations conducted with SLiM (Haller and Messer, 2019), and numerically with fastDTWF (Spence *et al*., 2023).

Using these transition probabilities, we computed the expected frequency in the descendant population conditional on the ancestral frequency. Results for our equilibrium model are presented in Figure 2D, with expected allele frequencies remaining unchanged over time under a neutral model; increasing under positive selection; decreasing under negative selection; and behaving in a frequency-dependent manner under stabilizing selection. For non-equilibrium models, the results are qualitatively consistent (Figure S1), but by shrinking population sizes, bottlenecks increase the effect of genetic drift, thereby reducing the differences between selection and neutrality (Figure S1A, S1C, S1D). Transition probabilities were concordant between SLiM simulations and fastDTWF, though there were some discrepancies at low allele frequencies due to the different assumptions of both methods (Methods, Figure S1A-C).

We next considered the impact of selection and demography backwards in time. In other words, given an allele segregating in a descendant population, what do we expect its frequency to have been in an ancestral population? Under equilibrium demography, we found that selected alleles are expected to be at lower frequencies in an ancestral population regardless of the direction of selection. Moreover, conditional on the frequency in the descendant population, the distribution of frequencies in the ancestral population is identical for positive and negative selection (Figure 3B), consistent with a classic result from Maruyama (1974).

**Figure 3:**
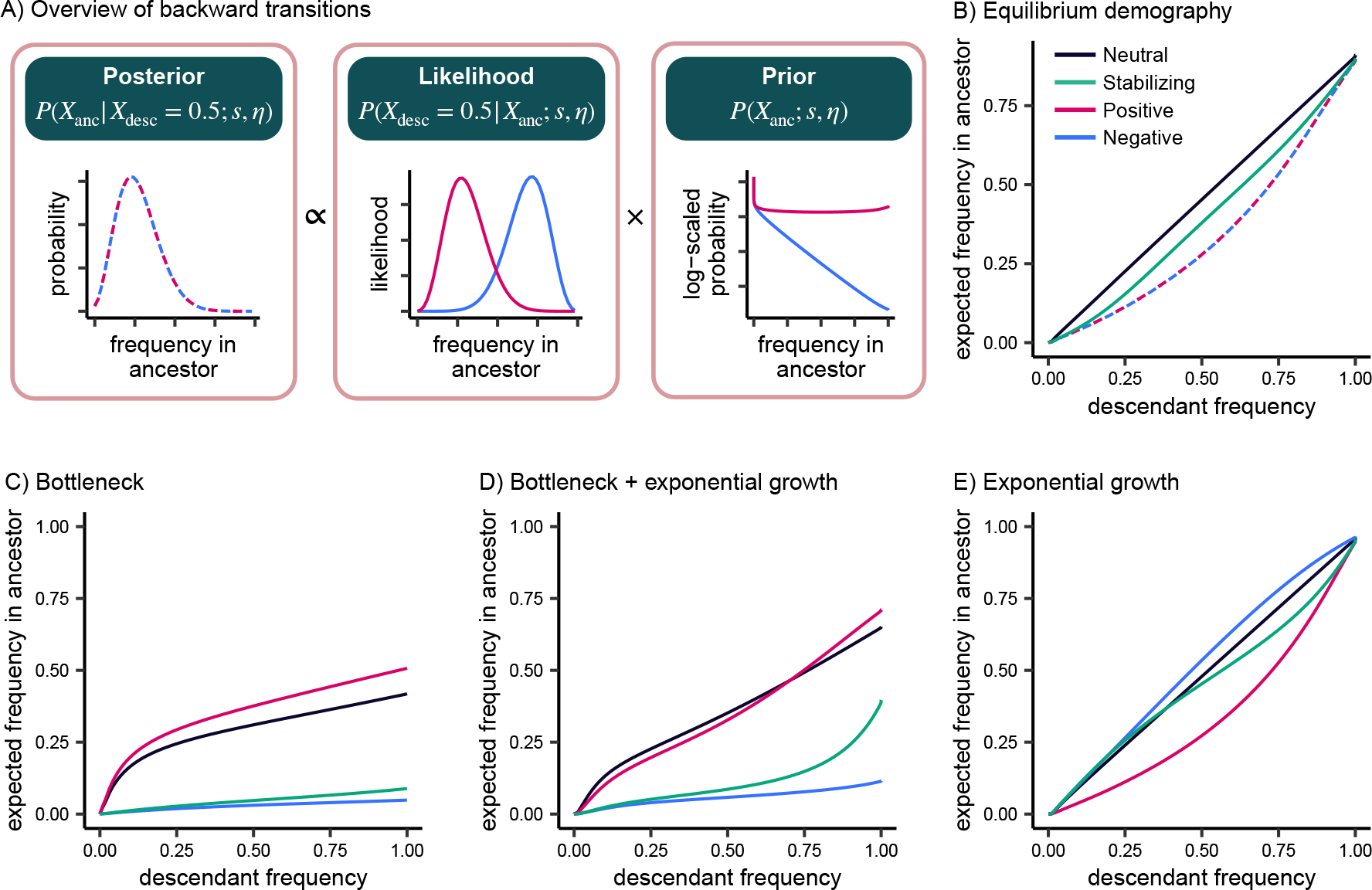
Backward transitions. **A)** Backward transition probabilities can be interpreted as a posterior consisting of a likelihood (i.e., forward transition probabilities), multiplied by a prior (i.e., the marginal distribution in the ancestor). **B-E)** Expected frequency in an ancestral population conditional on the frequency in the descendant population, computed with fastDTWF. Selection coefficients correspond to |*hs*| = 5.0 × 10*^−^*^4^. Demographic models consist of an ancestral population with *N_e_* = 10,000 and **B)** constant population size for 2,000 generations; **C)** 0.1*N_e_* bottleneck; **D)** 0.1 *N_e_* bottleneck and exponential growth at a rate of 0.1%; **E)** exponential growth at a rate of 0.1% each generation. To enhance visibility, overlapping distributions are represented with dashed lines.

This can be rationalized by thinking of the backward transition probability *P* (*X*_anc_ = *x*_anc_|*X*_desc_ = *x*_desc_; *s, η*) as a posterior probability proportional to the likelihood of transitioning from some ancestral frequency to the descendant frequency, *P* (*X*_desc_ = *x*_desc_|*X*_anc_ = *x*_anc_; *s, η*), weighted by the prior probability of being at that ancestral frequency, *P* (*X*_anc_ = *x*_anc_; *s, η*) (Figure 3A; Equation 2). Under equilibrium demography and negative selection, the likelihood is maximized by values of *x*_anc_ greater than *x*_desc_ because negative selection decreases frequencies over time. Conversely, under positive selection, the likelihood is maximized by values of *x*_anc_ less than *x*_desc_ because positive selection increases frequencies over time. However, the prior distribution differs under negative and positive selection in the opposite way: high derived allele frequencies in the ancestor are much less probable under negative selection compared to positive selection. For equilibrium demographies, these two forces exactly balance, resulting in negative and positive selection having identical posterior distributions (Figure 3A).

Under non-equilibrium demography, different trends emerged (Figure S2). The demographic models we considered only differ from each other following the split from the ancestral population; thus, while selection impacts both the prior *P* (*X*_anc_ = *x*_anc_; *s, η*) and the likelihood *P* (*X*_desc_ = *x*_desc_|*X*_anc_ = *x*_anc_; *s, η*), our demographic models only change the likelihood.

In the presence of a bottleneck, the increased strength of genetic drift makes the likelihood flatter, and as a result, the posterior is dominated by the prior, such that the expected frequency in the ancestor does not vary dramatically across descendant frequencies (Figure 3C, S2C).

Specifically, we found that alleles subject to negative and stabilizing selection are likely to be rare in the ancestor regardless of their frequency in the present. We found that a bottleneck followed by exponential growth reverses some of the observed flattening because the increase in population size decreases the strength of genetic drift (Figure 3D, S2D).

Under exponential growth alone, genetic drift is much weaker, meaning the likelihood is sharper than it is at equilibrium, and the posterior is dominated by the likelihood instead of the prior. As a result, alleles subject to negative selection are expected to be at higher frequencies in the ancestral population relative to alleles evolving neutrally, while alleles subject to positive selection are expected to be at lower frequencies (Figure 3E, S2E). We also varied the strength of the bottleneck and the degree of exponential growth, finding qualitatively similar results (Figure S3).

### Selection and the out-of-Africa demographic model

Having developed intuition for forward and backward allelic dynamics under simple demographies, we next considered their implications for realistic human demographies. We generated allele frequency distributions under a demographic model inferred from Han Chinese in Beijing, China (CHB); Yoruba in Ibadan, Nigeria (YRI); and Northern Europeans from Utah (CEU) in the 1000 Genomes Project (Figure 4A) (Auton *et al*., 2015; Jouganous *et al*., 2017). Briefly, this model consists of an ancestral human population under equilibrium demography with 24,000 individuals approximately 4,100 generations ago. At this time, the out-of-Africa migration occurs, and the population of the out-of-Africa branch shrinks by almost 90%, reflecting a severe bottleneck. After an additional 2,500 generations, the out-of-Africa branch splits into the CEU and CHB populations, both of which undergo a modest bottleneck and then experience exponential growth until the present day.

**Figure 4:**
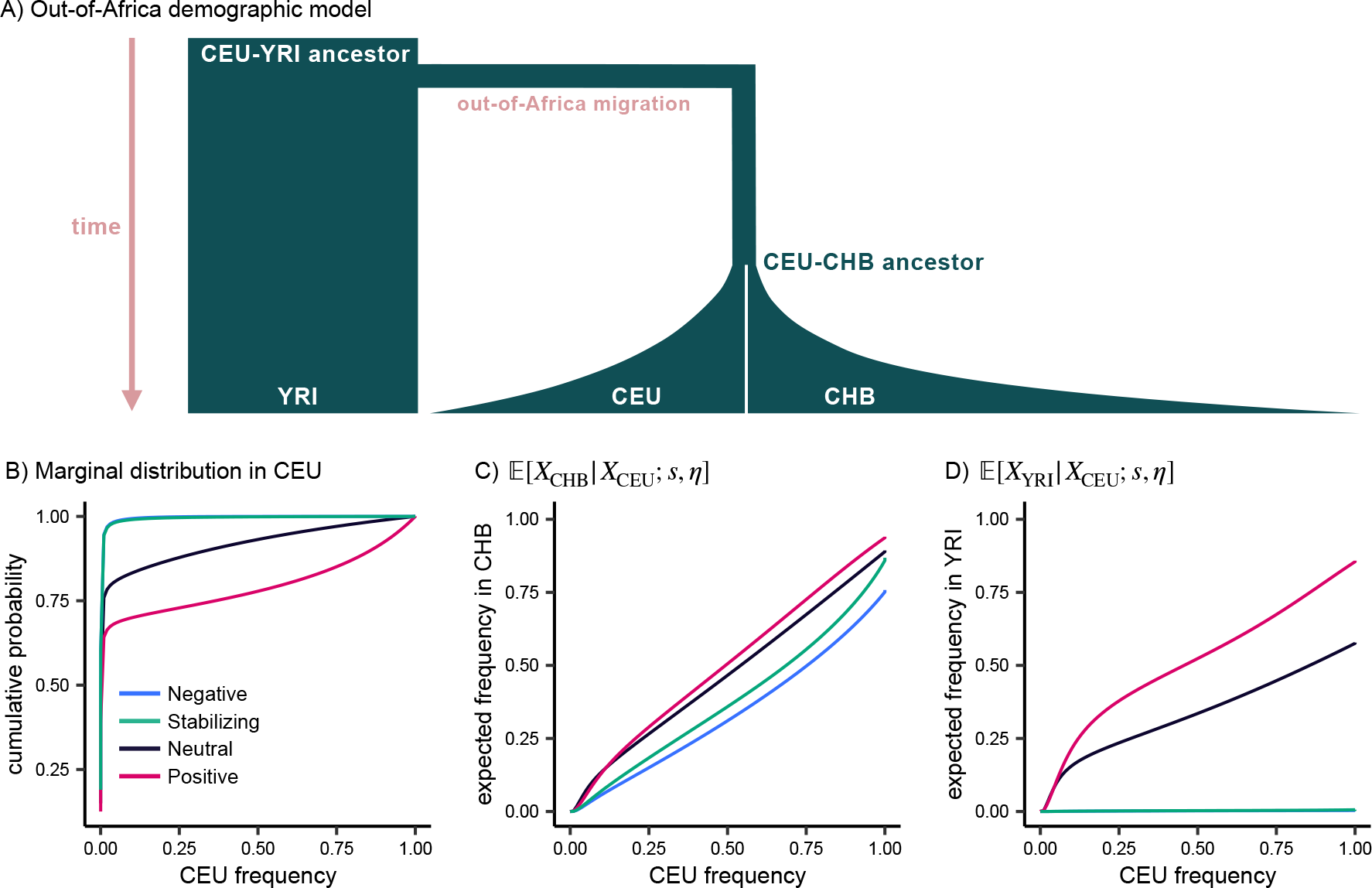
Out-of-Africa demography. **A)** Overview of out-of-Africa demographic model inferred from YRI, CEU, and CHB by Jouganous *et al*. (2017). Widths and lengths of branches are approximately proportional to population sizes and split times. **B)** Marginal distribution in CEU predicted by our theoretical results. **C, D)** Expected frequency in CHB and YRI conditional on the frequency in CEU. For all panels, selection coefficients correspond to *hs* = 5.0 10*^−^*^4^, computed with fast-DTWF.

We first sought to understand the impact of selection on frequency spectra in ancestral populations conditioned on the present-day frequency in CEU. Surprisingly, we found that the expected frequency in the ancestor of CEU and CHB, 𝔼[*X*_CEU_*_−_*_CHB ancestor_|*X*_CEU_; *s, η*], is similar for alleles evolving neutrally and those under strong negative selection (*s* = 10*^−^*^3^) (Figure S4A).

One might suppose that selection resembles neutrality because the 1,600 generations separating the CEU-CHB ancestor from CEU does not provide enough time for selection to act. However, this explanation can be ruled out by considering allele frequency dynamics forwards in time along this branch: it is apparent that selection substantially impacts allele frequencies over the 1,600 generations between the CEU-CHB ancestor and CEU (Figure S4B). Instead, our observations can be explained by non-equilibrium demography, which results in offsetting effects between the likelihood of CEU frequencies and the prior distribution in the CEU-CHB ancestor, similar to what we saw previously (Figure 3A).

In contrast, we found that the expected frequency in the ancestor of CEU and YRI, 𝔼[*X*_CEU_*_−_*_YRI ancestor_|*X*_CEU_; *s, η*], is similar for alleles under strong stabilizing selection and alleles under strong negative selection (Figure S4C). Regardless of the frequency in CEU, alleles under strong stabilizing or negative selection are likely to be maintained at low frequencies in the ancestor of CEU and YRI. This is concordant with the results we saw for a pure bottleneck scenario (Figure 3C), suggesting that the strength of the out-of-Africa bottleneck outweighs the subsequent exponential growth.

The stark difference in the backward transition probabilities for the CEU-YRI ancestor and the CEU-CHB ancestor is particularly notable given the qualitative similarity of the forward transition probabilities (Figure S4B, S4D). Ultimately, this underscores our finding that backward transitions are more sensitive to demography than forward transitions.

We next turned to understanding the allele frequency spectrum in one modern human population conditional on the frequency in another modern population. We focused on the distribution of frequencies in YRI and CHB conditional on the frequency in CEU, given that the majority of GWAS participants are highly genetically similar to CEU (Mills and Rahal, 2019).

We observed that conditional on the frequency in CEU, stabilizing and negative selection both result in lower frequencies in CHB and YRI relative to neutrality, regardless of the CEU frequency (Figure 4C, 4D). At low frequencies in CEU, positive selection and neutrality result in similar expected frequencies in CHB and YRI. However, as the CEU frequency increases, positively selected alleles are expected to be at much higher frequencies in CHB and YRI relative to neutral alleles. To understand this result, note that low frequency alleles tend to be younger on average and less likely to be segregating in the ancestral populations, especially when subject to selection (Figure S5A, S5D) (Maruyama, 1974). As the CEU frequency increases, the likelihood of a positively selected allele segregating in the ancestral population increases, such that the allele can experience positive selection along the other branches, resulting in higher frequencies in CHB and YRI relative to neutrality (Figure S5C, S5F).

Finally, we considered which conditional frequency spectra would be most informative for identifying the mode of selection acting on complex traits. We found that comparing YRI and CEU is more informative than comparing CHB and CEU, especially when selection is weak (Figure S6). We also found that it is generally much easier to distinguish neutrality from modes of selection than it is to distinguish *among* the different modes of selection. Specifically, frequency spectra in YRI conditional on CEU are similar under strong negative selection and strong stabilizing selection. At high (derived) allele frequencies in CEU, stabilizing selection and negative selection become easier to distinguish (Figure S5), but alleles at such high frequencies are discovered infrequently, particularly under these modes of selection (Figure 1A, 4B). Interestingly, stabilizing and negative selection are easier to distinguish when selection is weaker — in this regime, derived alleles are more likely to reach frequencies greater than 0.5, where the two modes of selection exhibit different allelic dynamics (Figure S7A-C, 2C).

### Empirical analysis of trait-associated variants

To understand the mode of selection acting on human complex traits, we investigated empirical conditional frequency spectra. There are two primary challenges in using our theoretical work on conditional frequency spectra to interpret empirical data. First, as we observed in our theoretical work, conditional frequency spectra are sensitive to demography, meaning that our quantitative theoretical results are sensitive to the exact demographic parameters in the Jouganous *et al*. (2017) out-of-Africa model. However, our qualitative theoretical results are generally robust to small changes in demographic parameters (Figure S8). Second, we expect trait-associated variants ascertained with GWAS to represent a mixture of different selection coefficients. These challenges make it difficult to draw quantitative conclusions from conditional frequency spectra for trait-associated variants. Instead, we qualitatively investigate the mode of selection acting on complex traits by comparing conditional frequency spectra for trait-associated variants to conditional frequency spectra for “matched variants”, a set of putatively neutral, non-associated variants with similar genomic properties.

We generated a set of matched variants by matching trait-associated variants to non-associated variants on two metrics: derived allele frequency in the UK Biobank White British cohort, and B-value, a measure of background selection (Murphy *et al*., 2022). Matching on B-value accounts for differences in allele frequency spectra that are due to selection at linked sites, as opposed to selection at the focal variant. Though some fraction of these matched variants may be functional and under selection, this only makes our comparisons more conservative.

We compared conditional frequency spectra in YRI and CHB between trait-associated variants and matched variants. We conditioned on frequencies in the UK Biobank White British cohort because it is the GWAS cohort for 94 of the 106 complex traits we analyzed, and because the individuals in this cohort are generally genetically similar to the individuals in the remaining GWAS cohorts. To compare conditional frequency spectra between trait-associated variants and matched variants, we first grouped variants into 10 deciles based on their frequency in the UK Biobank White British. Within each decile, we computed the mean frequency in YRI and CHB for trait-associated variants and matched variants.

Across all deciles, we found that variants associated with quantitative traits have a systematically lower mean frequency in YRI relative to matched variants (Figure 5A; p = 2.4 × 10*^−^*^93^; unpaired two-sided t-test). The same pattern was broadly observed for CHB, albeit to a lesser extent (Figure 5B; p = 9.8 × 10*^−^*^22^; unpaired two-sided t-test). For both YRI and CHB, much of this signal is driven by more trait-associated variants being lost in these populations relative to matched variants (Figure S9, S10).

**Figure 5:**
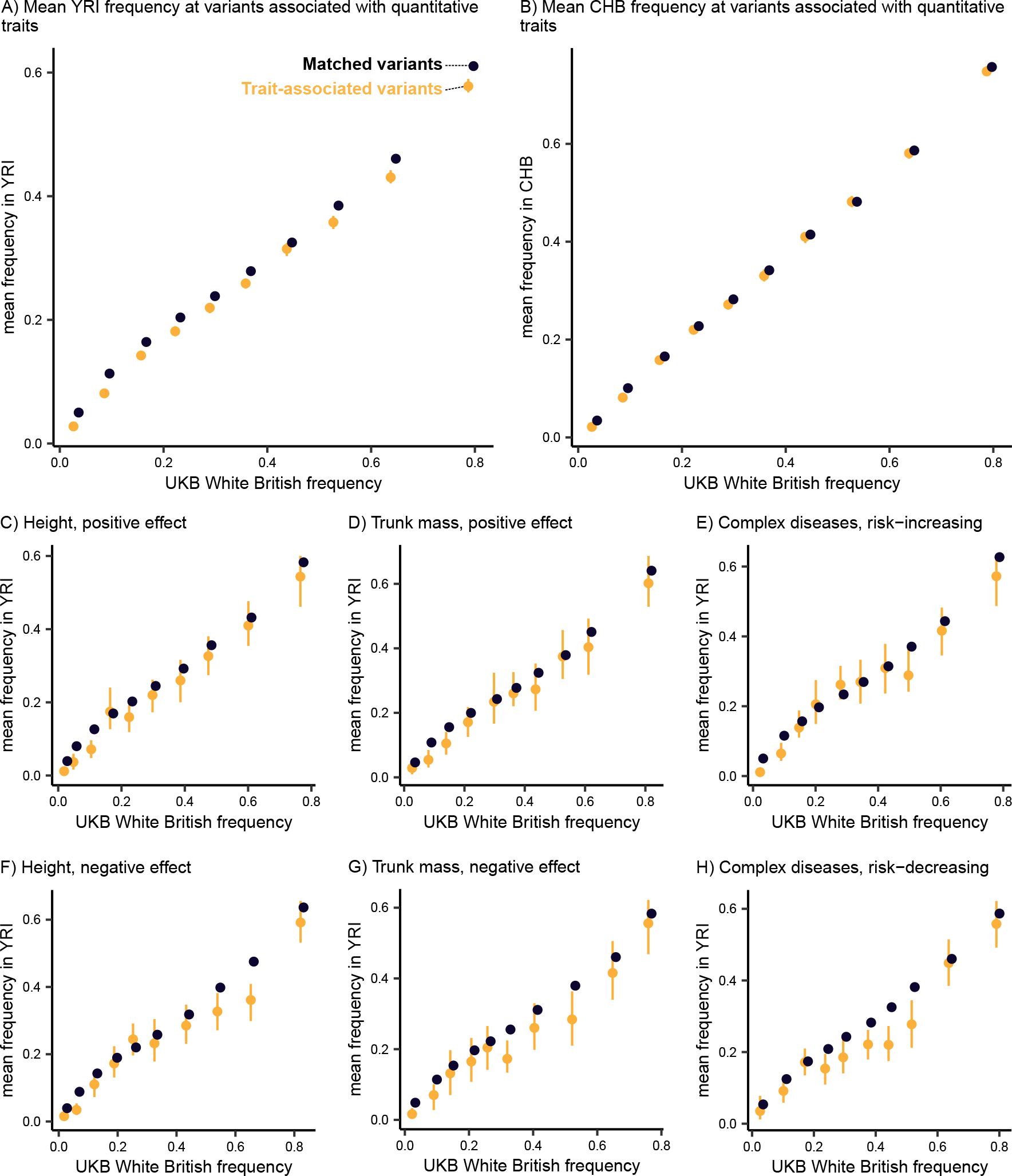
Conditional frequency spectra for trait-associated variants. **A, B)** Mean frequency in YRI and CHB conditional on UK Biobank White British frequency decile for quantitative trait-associated variants and matched variants. **C-H)** Mean frequency in YRI conditional on UK Biobank White British frequency decile for variants associated with height, trunk mass, and complex diseases. For all panels, error bars depict the 95% confidence interval for the mean, calculated from 100 bootstrap samples. Points are jittered along the x-axis (UK Biobank White British frequency) for better visibility.

Having established that trait-associated variants appear non-neutral, we aimed to understand which mode of selection explains the signal. We considered three models of selection on complex traits: stabilizing selection, directional selection, and purifying selection (i.e. negative selection against new mutations). Based on our theoretical results, stabilizing and negative selection on alleles produce similar conditional frequency spectra (Figure 4D), making it difficult to distinguish between stabilizing and purifying selection at the trait level. However, these two models are easy to distinguish from directional selection: if a trait is under directional selection, then alleles with positive and negative effects on the trait should experience directional selection in opposite directions, and hence have different conditional frequency spectra. This means that under directional selection, *either* trait-associated variants with a positive effect *or* trait-associated variants with a negative effect should be at higher frequencies in YRI and CHB relative to matched variants. Under stabilizing or purifying selection, however, we always expect to see trait-associated variants at lower frequencies in YRI and CHB relative to matched variants, regardless of the sign of their effect.

For each trait, we divided trait-associated variants into two groups based on the sign of their effect, and compared their conditional frequency spectra to matched variants. Specifically, we performed one-sided tests of the alternate hypothesis that trait-associated variants have lower frequencies in YRI relative to matched variants. Across quantitative traits, we found that trait-associated variants have significantly lower frequencies in YRI relative to matched variants, regardless of the sign of their effect (Figure 5C, 5D, 5F, 5G; p = 3.2 × 10*^−^*^10^ positive effect on height, p = 2.3 × 10*^−^*^8^ negative effect on height, p = 4.6 × 10*^−^*^4^ positive effect on trunk mass, p = 7.6 × 10*^−^*^3^ negative effect on trunk mass; unpaired one-sided t-test; see also Figure S11, S12). For complex diseases, we also found that trait-associated variants have lower frequencies in YRI relative to matched variants, regardless of whether they are risk-increasing or risk-decreasing (Figure 5E, 5H; p = 1.4 × 10*^−^*^6^ risk-increasing, p = 1.5 × 10*^−^*^11^ risk-decreasing; unpaired one-sided t-test). Thus, we do not find evidence for directional selection, and instead find compelling evidence that stabilizing or purifying selection is the predominant mode of selection acting on complex traits and diseases.

Given that our results depend on knowing which allele is ancestral or derived, one consideration is the potential mispolarization of variants. In particular, variants at CpG sites are more difficult to polarize because of the higher mutation rate at these sites (Jónsson *et al*., 2017; Keightley and Jackson, 2018). To account for this, we repeated our analyses excluding all transition mutations, and found that our results were qualitatively unchanged (Figure S13).

Another potential consideration is that our set of trait-associated variants likely contains some non-functional “tag” variants in linkage disequilibrium (LD) with the true causal variants: if two variants are in LD, their association signals are also highly correlated, making it difficult to determine which variant is causal. Ascertainment of non-functional tag variants can affect our results in two ways. First, tag variants and causal variants could be at different frequencies, distorting our empirical conditional frequency spectra. Second, conditional frequency spectra encode information about *derived* allele frequencies. If the derived allele at the tag variant is in negative LD with the derived allele at the causal variant, the trajectory of the derived allele at the tag variant will be opposite of that at the causal variant. Then, in conditional frequency spectra, the true direction of selection would appear to be reversed. While this would make our qualitative test of neutrality against selection more conservative, it could bias our results when trying to determine the mode of selection.

We used coalescent simulations to better understand these potential sources of bias. First, we investigated whether variants in LD could be at dramatically different frequencies. We found that for variants with *r*^2^ ≥ 0.5, the corresponding minor allele frequencies are essentially identical with high probability (Figure S14A). Next, we estimated the probability that the derived allele at a tag variant is in negative LD with the derived allele at a causal variant. We found that negative LD is uncommon when the derived allele at the tag variant is rare, but negative LD between derived alleles happens with ∼50% probability when the derived allele at the tag variant reaches higher frequencies (Figure S14B).

Given that rare derived alleles more reliably tag derived causal variants, we repeated our tests for directional selection using trait-associated variants in the lowest quintile of UK Biobank White British derived allele frequency. For quantitative traits, we found that our results were qualitatively unchanged; we find that both trait-increasing and trait-decreasing variants have significantly lower frequencies in YRI relative to matched variants, ruling out directional selection (p = 2.4 × 10*^−^*^9^ positive effect on height, p = 1.2 × 10*^−^*^8^ negative effect on height, p = 5.7 × 10*^−^*^4^ positive effect on trunk mass, p = 7.4 × 10*^−^*^4^ negative effect on trunk mass; unpaired one-sided t-test). For complex diseases, we found that risk-increasing variants had significantly lower frequencies than matched variants but did not find significant evidence for risk-decreasing variants (p = 1.7 × 10*^−^*^15^ risk-increasing, p = 0.09 risk-decreasing; unpaired one-sided t-test). This could be due to the reduction in power when excluding 80% of trait-associated variants, or could be compatible with directional selection against variants increasing disease risk.

### Impact of selection and demography on polygenic score portability

Our results have broader implications for applications of GWAS data, particularly in the context of polygenic scores. Polygenic scores estimate the genome-wide genetic contribution to a trait or disease using trait-associated variants ascertained in a GWAS. The phenotypic variance explained by polygenic scores is known to decrease in populations with less genetic similarity to the GWAS cohort — commonly referred to as a lack of “portability” (Ding *et al*., 2023; Martin *et al*., 2019). The portability of a polygenic score is influenced by many factors including differing patterns of linkage between ascertained variants and causal variants across populations; bias in effect sizes due to population structure; and environmental differences across populations (Mostafavi *et al*., 2020; Patel *et al*., 2022; Privé *et al*., 2022; Wang *et al*., 2020; Wojcik *et al*., 2019; Vilhjálmsson *et al*., 2015). Here, we focus on understanding the impact of allele frequency differences on portability, particularly in the context of different modes of selection.

For quantitative traits, the phenotypic variance explained by a polygenic score is proportional to 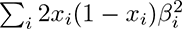: the sum of squared effect sizes, 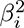, at trait-associated variants, weighted by heterozygosity, 2*x_i_*(1 − *x_i_*). Thus, systematic differences in heterozygosity across populations will drive differences in the variance explained by the polygenic score, affecting portability.

Using theoretical conditional frequency spectra, we computed the expected heterozygosity of variants in CHB and YRI conditional on their frequency in CEU. We found that in both CHB and YRI, stabilizing and negative selection reduce heterozygosity relative to neutral alleles for low to moderate CEU frequencies (Figure 6A, 6B). At CEU frequencies greater than 0.75, however, we found that negative and stabilizing selection can actually increase heterozygosity in CHB above neutral levels.

**Figure 6:**
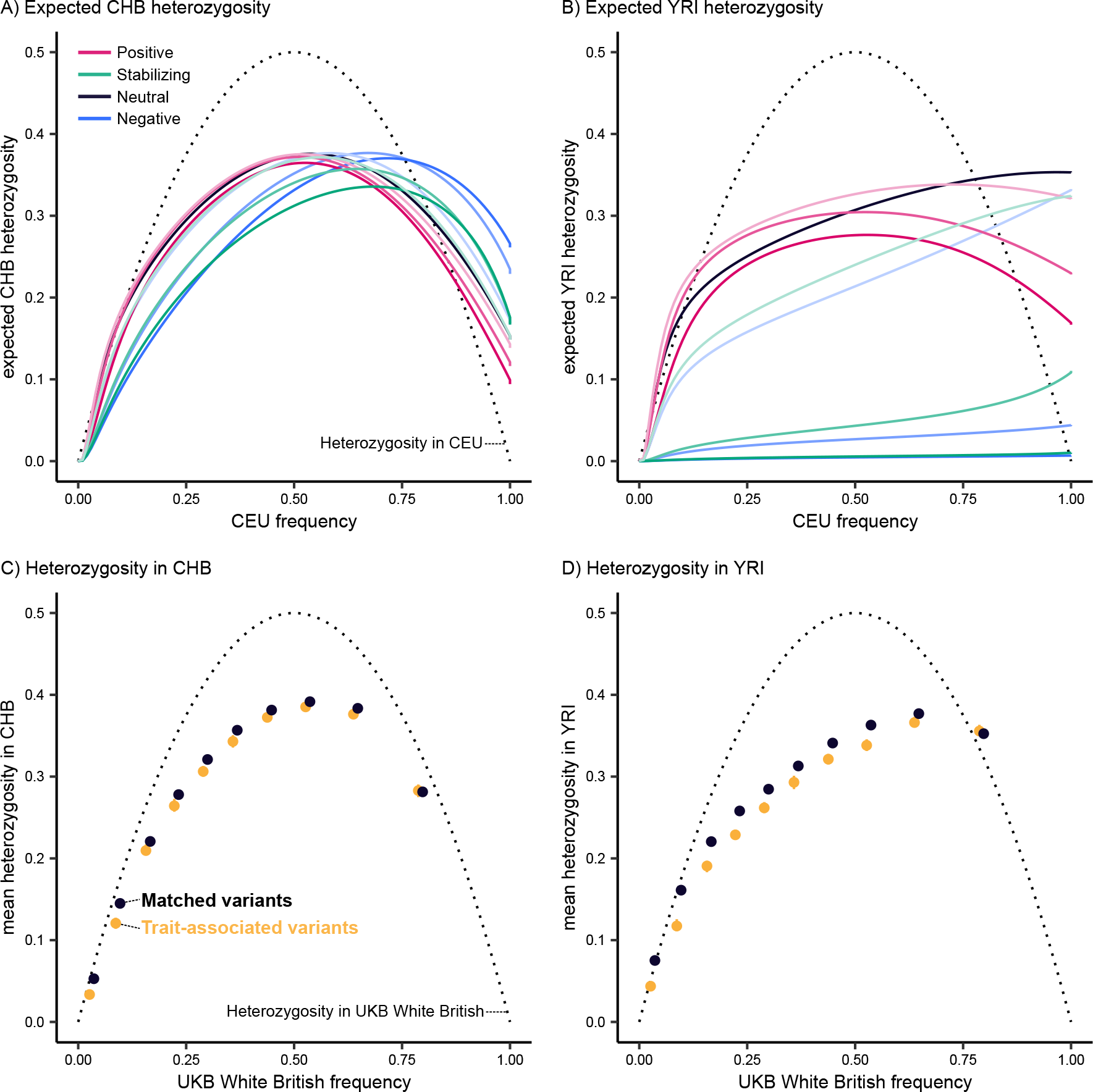
Implications for polygenic score portability. **A, B)** Expected heterozygosity in CHB and YRI conditional on the frequency in CEU, computed with fastDTWF. Selection coefficients range from *hs* = 5.0 10*^−^*^5^, shown in the lightest shades, to *hs* = 5.0 10*^−^*^4^, shown in the darkest shades. **C, D)** Mean heterozygosity in CHB and YRI conditional on UK Biobank White British frequency decile for all trait-associated variants and matched variants. Error bars depict the 95% confidence interval for the mean, calculated from 100 bootstrap samples, and points are jittered along the x-axis (UK Biobank White British frequency) for better visibility. For all panels, dotted line corresponds to the heterozygosity in the conditional population.

Relative to CHB, selection has a much more dramatic impact on the heterozygosity of alleles in YRI (Figure 6A, 6B). Regardless of CEU frequency, negative and stabilizing selection result in extremely low heterozygosity in YRI. Even weakly selected alleles (corresponding to *Ns* ≈ 1) have a substantially reduced heterozygosity in YRI, almost 10% lower than neutral alleles. In CHB, however, the heterozygosity under weak selection is nearly indistinguishable from neutrality.

This suggests that selection acting on complex traits (or correlated traits) will strongly impact polygenic score portability from CEU-like populations to YRI-like populations. To understand the role of demography, we considered a simpler demographic model consisting of two populations that split 2,000 generations ago and maintained a constant population size of *N_e_* = 10,000. We computed the expected heterozygosity in one population conditional on the other and again found that selected alleles have reduced heterozygosity in comparison to neutral alleles (Figure S15). However, the magnitude of this difference is much smaller than what our theoretical results predict for CEU and YRI. It is therefore evident that while selection reduces polygenic score portability across all demographic scenarios, certain demographic scenarios can exacerbate the impacts of selection. Indeed, much of the reduction in polygenic score portability observed in empirical data can likely be attributed to the out-of-Africa bottleneck experienced by CEU-like populations (Martin *et al*., 2019; Ding *et al*., 2023).

To understand this phenomenon in empirical data, we compared the mean heterozygosity of trait-associated variants and matched variants in CHB and YRI across each decile of UK Biobank White British frequency. We found that trait-associated variants systematically have less heterozygosity in CHB and YRI, relative to matched variants (Figure 6C, 6D; p = 9.9 × 10*^−^*^39^ CHB, p = 5.0 × 10*^−^*^123^ YRI; unpaired two-sided t-test). As suggested by our theoretical results, the reduction in heterozygosity at trait-associated variants is stronger in YRI compared to CHB. These results illustrate how differences in allele frequencies driven by selection can contribute to the poor portability of polygenic scores.

## Discussion

We presented conditional frequency spectra as a tool for studying selection on complex traits. The utility of conditional frequency spectra stems from the peculiarities of GWAS: GWAS ascertainment implicitly prioritizes variants with large minor allele frequencies and large effect sizes, both of which are related to the strength of selection on a variant. In doing so, GWAS ascertainment obscures much of the information one actually wants to learn from GWAS. Here, we used conditional frequency spectra to circumvent the issues caused by GWAS ascertainment to study selection on complex traits — but conditional frequency spectra should be broadly useful for other statistical genetics applications as well.

We note that inferring selection coefficients from conditional frequency spectra is challenging. Though different models of selection are easily distinguished looking forwards in time, we find that backward transitions are sensitive to demography. This means that demographic misspecification can hinder the inference of selection coefficients from conditional frequency spectra. To account for this, one could use putatively neutral variation to infer a demographic model relating the specific cohorts of individuals represented in empirical conditional frequency spectra — in our case, UK Biobank White British, 1000 Genomes CHB, and 1000 Genomes YRI. This demographic model could then be used to generate theoretical expectations of conditional frequency spectra and ultimately obtain more robust estimates of selection coefficients from trait-associated variants. In any case, care would need to be taken such that errors in the inferred demographic model do not affect estimates of selection.

Nevertheless, our theoretical analyses of conditional frequency spectra identified strong qualitative signatures for each model of selection. Applying this intuition to empirical data, we found significant evidence for stabilizing or purifying selection acting on trait-associated variants, but no evidence for directional selection. We note that our approach only enables us to detect sustained directional selection shared by all branches in the tree. Others have previously proposed that selection acts in divergent directions across human populations, structuring many complex trait phenotypes (Guo *et al*., 2018); our analyses cannot rule out this possibility. However, our work does highlight that certain empirical observations can be compatible with multiple models of selection on complex traits. This emphasizes the benefit of invoking explicit population genetic models: models are crucial for interpreting what observations can or cannot tell us about selection, even if inference under such models is difficult.

Lastly, we examined the consequences of selection on complex traits for polygenic score portability. Though it is well-known that selection on complex traits can affect the portability of polygenic scores (Durvasula and Lohmueller, 2021; Wang *et al*., 2020; Yair and Coop, 2022), our approach enables us to contrast different models of selection, as well as understand the respective contributions of demography and selection. We find that while selection reduces portability, the out-of-Africa bottleneck and subsequent drift likely plays an outsized role in the decreased portability from CEU-like populations to YRI-like populations. Here, we only modeled alleles, as opposed to traits, and so our results cannot speak to the effects of environmental heterogeneity, different contributions of the additive genetic component across populations, gene-gene interactions, or gene-environment interactions. Investigating these aspects further is likely to be informative for improving polygenic score portability across groups.

In conclusion, we characterize the conditional frequency spectrum under different models of selection, providing insight into properties of trait-associated variants and ultimately underscoring the value of conditioning on GWAS ascertainment.

## Methods

### Theoretical analysis of allele frequency spectra

We characterized the impact of demography and selection on the distribution of allele frequencies across populations by treating demography, *η*, and selection, *s*, as fixed quantities in a discrete probability distribution over allele frequencies *X*. We considered four distinct types of allele frequency distributions:

i. Forward transitions *P* (*X*_desc_ = *x*_desc_|*X*_anc_ = *x*_anc_; *s, η*): the distribution of frequencies in a descendant population, conditional on the frequency in its ancestral population
ii. Marginals *P* (*X_k_* = *x_k_*; *s, η*): the marginal distribution of frequencies in population *k*
iii. Backward transitions *P* (*X*_anc_ = *x*_anc_|*X*_desc_ = *x*_desc_; *s, η*): the distribution of frequencies in an ancestral population, conditional on the frequency in its descendant population
iv. Conditional frequency spectra *P* (*X_j_* = *x_j_* |*X_k_* = *x_k_*; *s, η*): the distribution of frequencies *X* in population *j*, conditional on the frequencies in population *k*, where *j* and *k* share a common ancestor.

For each branch in a particular demographic model, we obtained forward transitions *P* (*X*_desc_ = *x*_desc_|*X*_anc_ = *x*_anc_; *s, η*) using two different implementations of the Wright-Fisher model. First, we used fastDTWF to numerically compute likelihoods under the discrete-time Wright-Fisher model (Spence *et al*., 2023). fastDTWF is restricted to modeling demographies that are piecewise constant. As such, we approximated exponential population growth by a piecewise constant model by updating the population size every 20 generations. Second, we used forward-time SLiM simulations, which are capable of modeling more flexible demographies, to obtain Monte Carlo approximations of the forward transitions (Haller and Messer, 2019). We started at an initial ancestral frequency *x*_anc_ and simulated forward in time, recording the frequency in every downstream population in the demographic model. We then approximated the forward transition *P* (*X*_desc_ = *x*_desc_|*X*_anc_ = *x*_anc_; *s, η*) with the empirical distribution across replicate simulations. We performed 1,000 replicates for each initial frequency for simple two-population demographic models and 2,000 replicates for the out-of-Africa demographic model (see *Modeling demography* below for details). The forward transitions generated by fastDTWF and SLiM simulations differed in that while fastDTWF incorporates both recurrent mutations and new mutations, our particular implementation of SLiM simulations requires alleles to be segregating in the ancestral population of the demographic model.

To compute a marginal distribution for a descendant population, we first obtained the marginal distribution of the ancestral population in the demographic model, *P* (*X*_anc_; *s, η*), using fastDTWF. We then computed the marginal distribution for the descendant population as follows:

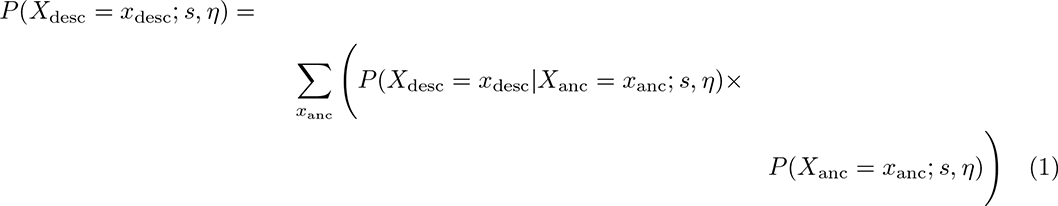

Using the marginal probability *P* (*X*_desc_ = *x*_desc_; *s, η*), we computed the backward transition as follows:

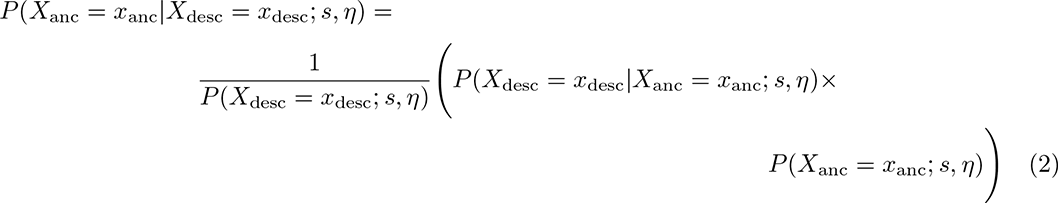

Similarly, we computed a conditional frequency distribution *P* (*X_j_* = *x_j_* |*X_k_* = *x_k_*; *s, η*) in which populations *j* and *k* share a common ancestor:

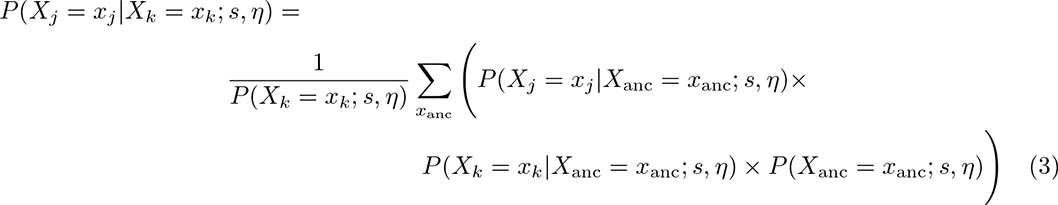

In practice, we performed these computations on frequency distributions by first aggregating allele frequencies into bins. A distribution over allele frequencies is a discrete probability distribution with 2*N* possible outcomes, where *N* is the diploid population size. Thus, without binning frequencies, the number of computations required for basic operations quickly becomes cumbersome. Moreover, adjacent conditional distributions will be trivially similar (e.g. the probability distribution *P* (*X_j_* |*X_k_* = 10000 counts; *s, η*) is nearly identical to the probability distribution *P* (*X_j_* |*X_k_* = 10001 counts; *s, η*)) (Spence *et al*., 2023). To account for the fact that adjacent conditional distributions are more dissimilar when alleles are close to loss or fixation, we used denser binning for allele frequencies close to 0 and 1. Specifically, we created bins corresponding to 0 derived allele counts, 1 count, 2 counts, (2, 5] counts, (5, 10] counts, and (10, 20] counts to cover the space of allele frequencies close to 0; and bins corresponding to 2*N* counts, 2*N* − 1 counts, and (2*N* − 5, 2*N* − 2] counts to cover the space of allele frequencies close to 1. To cover the remaining space of allele frequencies, we created frequency bins with a width of 0.01. For each population this procedure generated roughly 100 allele frequency bins, regardless of the population size *N*.

### Modeling demography

We relied on two different types of demographic models. First, we considered a simple two population model consisting of one ancestral population and one descendant population separated by 2,000 generations. We used this model to understand how selection impacts allele frequency dynamics under different demographic conditions: equilibrium demography (i.e. constant population size), a bottleneck (i.e. a sudden decrease in population size), exponential population growth, and a bottleneck followed by exponential growth. We focused on parameter values relevant to human demographic history, modeling an ancestral population size with *N_e_* = 10,000; a bottleneck of 0.1*N_e_* and 0.3*N_e_*; and an exponential growth rate of 0.05% and 0.1% (Gutenkunst *et al*., 2009; Jouganous *et al*., 2017; Ragsdale and Gravel, 2019).

The second demographic model we considered was the out-of-Africa model inferred by Jouganous *et al*. (2017). Jouganous *et al*. (2017) inferred demographic parameters relating three modern-day populations in the 1000 Genomes Project (Auton *et al*., 2015): Han Chinese in Beijing, China (CHB); Yoruba in Ibadan, Nigeria (YRI); and Northern Europeans from Utah (CEU). In brief, this model consists of a common ancestor for CHB, YRI, and CEU 4,100 generations ago, followed by a strong out-of-Africa bottleneck and subsequent split between CHB and CEU 1,600 generations ago. CHB experiences a modest bottleneck after splitting from CEU, and the CHB and CEU branches both experience exponential growth following their split (Figure 3A). Our model only differs from that of Jouganous *et al*. (2017) in that we did not model migration between branches; our probability computations assume that populations are independent after branching (see *Theoretical analysis of allele frequency spectra* above).

### Modeling selection

We considered three types of selection on alleles: negative genic selection, positive genic selection, and stabilizing selection. Using common notational convention, we can represent the fitness of the *AA* genotype as 1, the *Aa* genotype as 1 + *hs*, and the *aa* genotype as 1 + *s*, where *A* is the ancestral allele and *a* is the derived allele (Gillespie, 2004). Under negative genic selection, *h* = 0.5 and *s <* 0. We considered three values of *s*: −1 × 10*^−^*^4^, −5 × 10*^−^*^4^, and −1 × 10*^−^*^3^. These values range from extremely weak (corresponding to *Ns* ≈ 1 in the ancestral population) to the strongest selection coefficients inferred for trait-associated variants ascertained in complex trait GWAS (Simons *et al*., 2022). Under positive genic selection, *h* = 0.5 and *s >* 0. We modeled values of *s* ranging from +1 × 10*^−^*^4^ to +1 × 10*^−^*^3^, analogous to negative selection. Under stabilizing selection, allele frequency dynamics resemble a scenario where *hs <* 0 but *s* = 0, such that only heterozygotes experience a fitness cost (Robertson, 1956; Walsh and Lynch, 2018). We modeled values of *hs* ranging from −5 × 10*^−^*^4^ to −5 × 10*^−^*^5^, analogous to negative selection.

### Empirical analysis of trait-associated variants

We obtained trait-associated variants from published GWAS for 94 quantitative complex traits and 12 (binary) complex diseases. For quantitative traits, we obtained a set of 18,229 variants ascertained in the UK Biobank “White British” cohort with a minor allele frequency of at least (http://www.nealelab.is/uk-biobank). The White British cohort consists of approximately 337,000 unrelated individuals in the UK Biobank. We restricted our analyses to this subset of the UK Biobank because our approach relies on performing GWAS in a cohort that is relatively homogeneous and has reasonably high genetic similarity to one of the populations in our out-of-Africa demographic model. Moreover, the size of the White British cohort enables less noisy estimation of allele frequencies.

For disease traits, we obtained a total of 1,040 variants ascertained in cohorts with European ancestries using individual case/control GWAS (Table S1) (Aragam *et al*., 2022; Bellenguez *et al*., 2022; International IBD Genetics Consortium (IIBDGC) *et al*., 2013; De Lange *et al*., 2017; Demontis *et al*., 2023; Ishigaki *et al*., 2022; Michailidou *et al*., 2015; Mullins *et al*., 2021; Nalls *et al*., 2019; Pardiñas *et al*., 2018; Schumacher *et al*., 2018; Scott *et al*., 2017). Paralleling our quantitative trait analyses, we filtered out variants with a minor allele frequency less than 0.01 in the UK Biobank White British.

For each trait-associated variant, we generated a set of “matched variants” that share similar properties. We started with approximately 11 million (imputed) variants in the UK Biobank that are not found to be trait-associated in our GWAS datasets. For a given trait-associated variant, we identified matched variants by selecting for two different criteria. First, we matched on the derived allele frequency in the UK Biobank White British. To compute derived allele frequencies, we obtained the ancestral allele state inferred by Auton *et al*. (2015) and stored in the Ensembl variation database for all trait-associated variants and all other imputed variants in the UK Biobank (Martin *et al*., 2023). Briefly, Auton *et al*. (2015) identified ancestral allele states using a multiple genome alignment between human, chimp, orangutan, and rhesus macaque to infer ancestral sequences and annotate ancestral allele states. We matched on UK Biobank White British derived allele frequency by binning continuous frequencies into 100 evenly spaced bins between 0 and 1; two derived allele frequencies were deemed to match if they belonged to the same bin.

Second, we matched variants on their B-value, a background selection statistic (Murphy *et al*., 2022). Background selection impacts allele frequency spectra by decreasing genetic diversity, and varies throughout the genome as a function of recombination rate, gene density, and other genomic features. Thus, matching on B-values accounts for differences in allele frequency spectra between trait-associated variants and matched variants that arise due to varying levels of background selection throughout the genome. We matched on B-value by binning the B-values for the 11 million variants imputed by UKB into 15 quantiles; two B-values were deemed to match if they belonged to the same bin. We discarded trait-associated variants with fewer than 500 matched variants, generating an average of 11,547 matched variants for each trait-associated variant.

We generated empirical conditional frequency spectra for both trait-associated variants and matched variants. We considered two types of conditional frequency spectra: the distribution of frequencies in CHB (Han Chinese from Beijing, China) conditional on the frequency in UK Biobank White British, and the distribution of frequencies in YRI (Yoruba in Ibadan, Nigeria) conditional on the frequency in UK Biobank White British. To compute allele frequencies in CHB and YRI, we used 103 CHB individuals and 108 YRI individuals from 1000 Genomes Phase 3, respectively.

To pool information across the relatively small number of trait-associated variants, we split them into ten deciles based on their frequency in UK Biobank White British. Within each UK Biobank frequency decile, we computed the mean frequency in CHB and YRI across all trait-associated variants in the decile. This approximates the expectation over the conditional frequency spectrum, 𝔼[*X*_CHB_|*X*_UKB_] and 𝔼[*X*_YRI_|*X*_UKB_], for trait-associated variants. We next computed the analogous quantity for matched variants by aggregating across the matched variants for each trait-associated variant in a particular UK Biobank frequency decile. To compute the 95% confidence interval for the mean(s), we separately bootstrapped over trait-associated and matched variants, and reported the 0.025 and 0.975 quantiles across 100 replicates.

To test for selection, we compared trait-associated and matched variants by performing an unpaired two-sided t-test for unequal sample variances within each UK Biobank frequency decile. We combined p-values across deciles using Fisher’s method.

To specifically test for directional selection, we split trait-associated variants into two groups based on the sign of their effect (i.e. trait-increasing and trait-decreasing). For each group, we tested the alternative hypothesis that trait-associated variants have a lower frequency in YRI and CHB relative to matched variants by performing an unpaired one-sided t-test for unequal sample variances within each UK Biobank frequency decile. We again combined p-values across deciles using Fisher’s method. When the null hypothesis was rejected for both trait-increasing and trait-decreasing variants, we interpreted this as evidence against directional selection.

### Analysis of imperfect tagging

We aimed to understand the probability that a trait-associated variant accurately tags the “correct” allele at the true causal variant — in other words, the probability that derived alleles at a pair of linked variants are positively correlated. Using msprime (Baumdicker *et al*., 2022), we performed coalescent simulations for 100 diploid individuals. Specifically, we simulated 100,000 coalescent trees with exactly two mutations. For each tree, we calculated the linkage disequilibrium (LD) between the pair of mutations as the squared correlation coefficient of their genotypes. We then computed the probability that derived alleles are linked at pairs of mutations meeting various LD thresholds.

## Data availability statement

Conditional frequency spectra and other allele frequency distributions can be found at https://github.com/roshnipatel/conditional-frequency-spectra, along with code for performing computations on frequency distributions and analyzing empirical data.

## Acknowledgments

We thank Alyssa Lyn Fortier and Courtney Smith for valuable feedback on the manuscript, and members of the Pritchard lab for helpful conversations about this work. Research reported in this publication was supported by NIH grants R01HG011432 and R01HG008140.

## Supplemental Tables and Figures

**Table S1:**
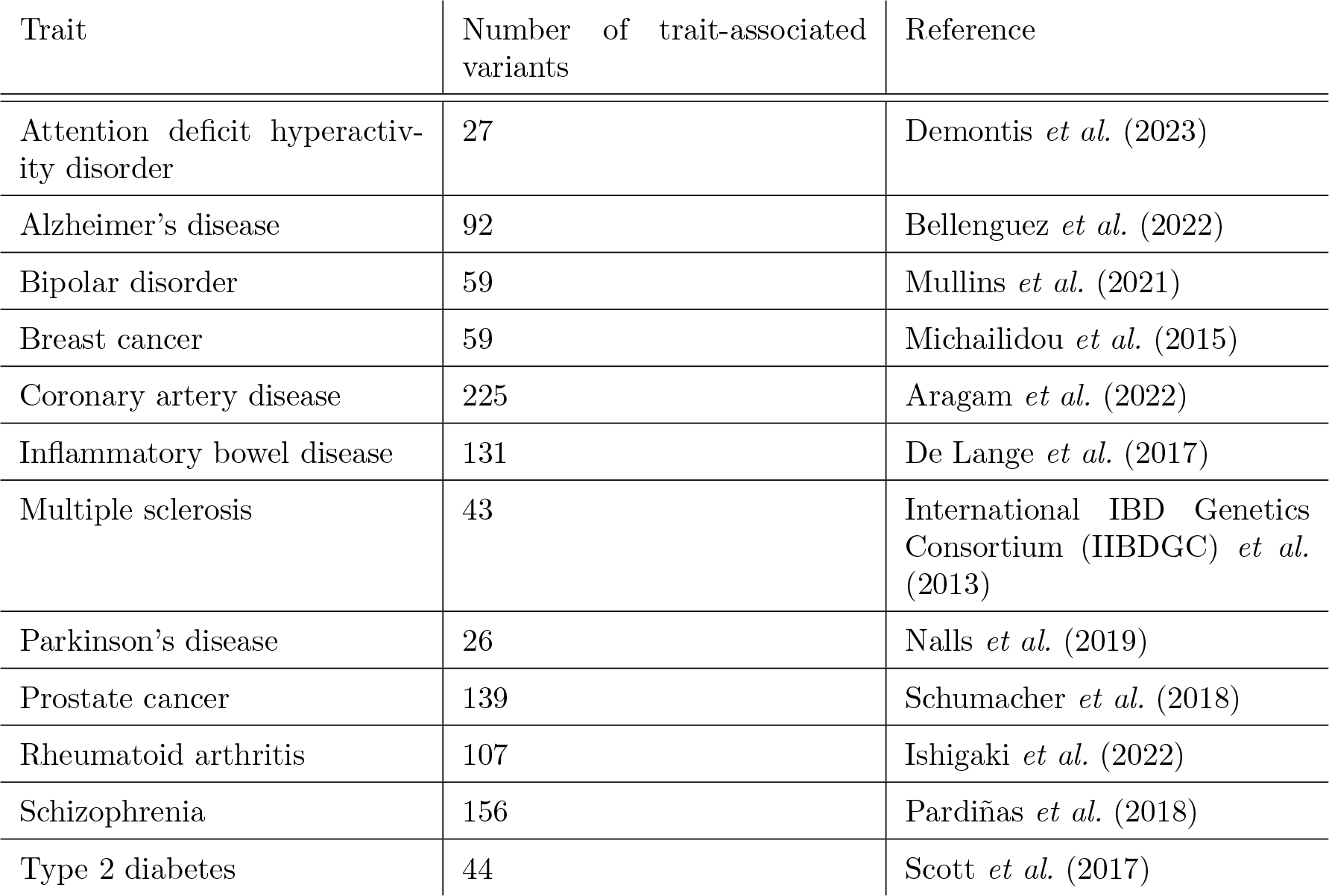
Details on complex disease GWAS.

**Figure S1:**
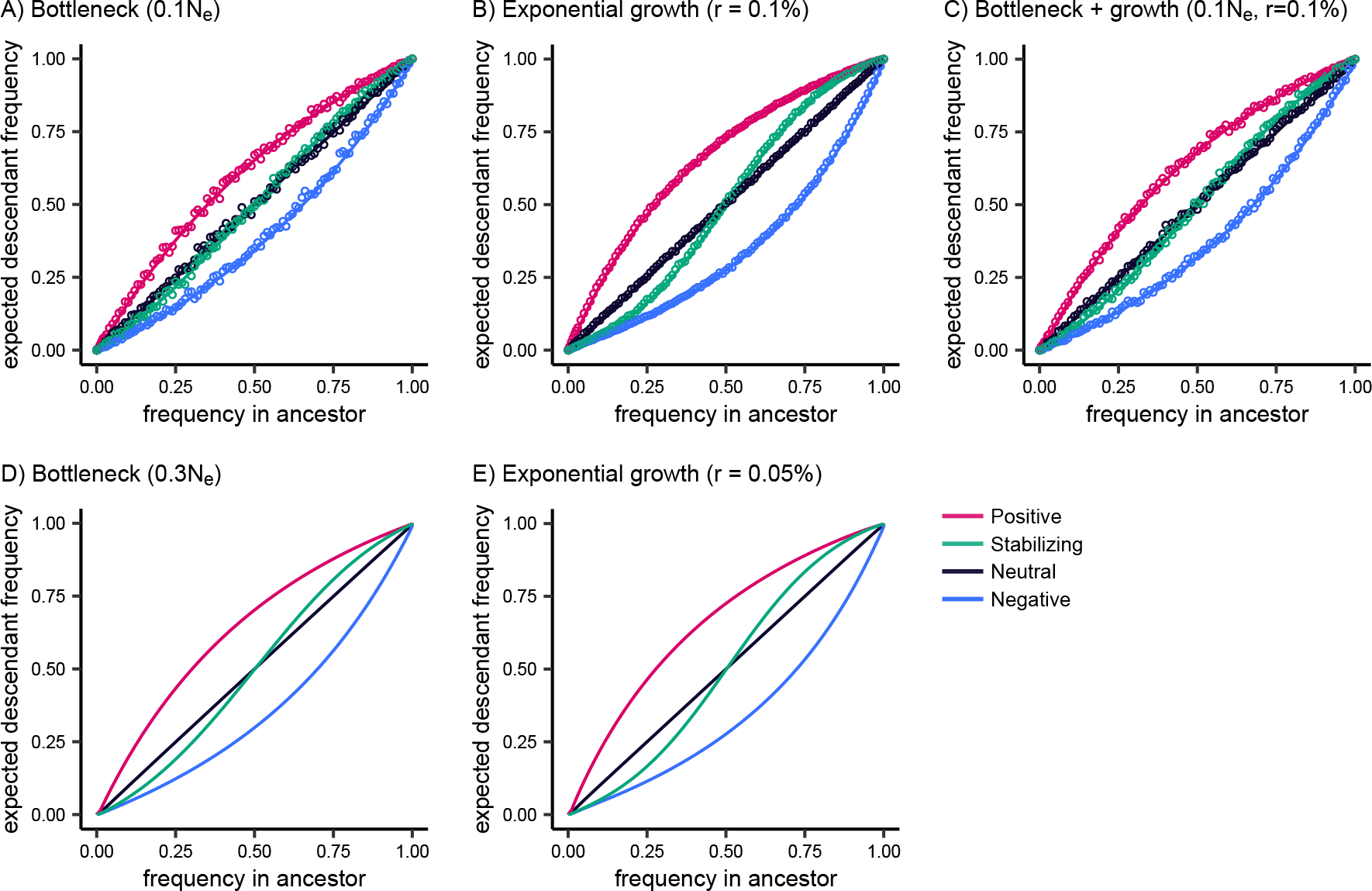
Forward transitions under different demographic models. Expected frequency in a descendant population conditional on the frequency in the ancestral population 2,000 generations prior, computed with fastDTWF (solid lines) and SLiM simulations (open circles). Selection coeffi-cients correspond to |*hs*| = 5.0 × 10*^−^*^4^. Demographic models consist of an ancestral population with *N_e_* = 10,000 and **A)** 0.1*N_e_* bottleneck; **B)** exponential growth at a rate of 0.1% each generation; **C)** 0.1*N_e_* bottleneck and exponential growth at a rate of 0.1%; **D)** 0.3*N_e_* bottleneck; **E)** exponential growth at a rate of 0.05% each generation.

**Figure S2:**
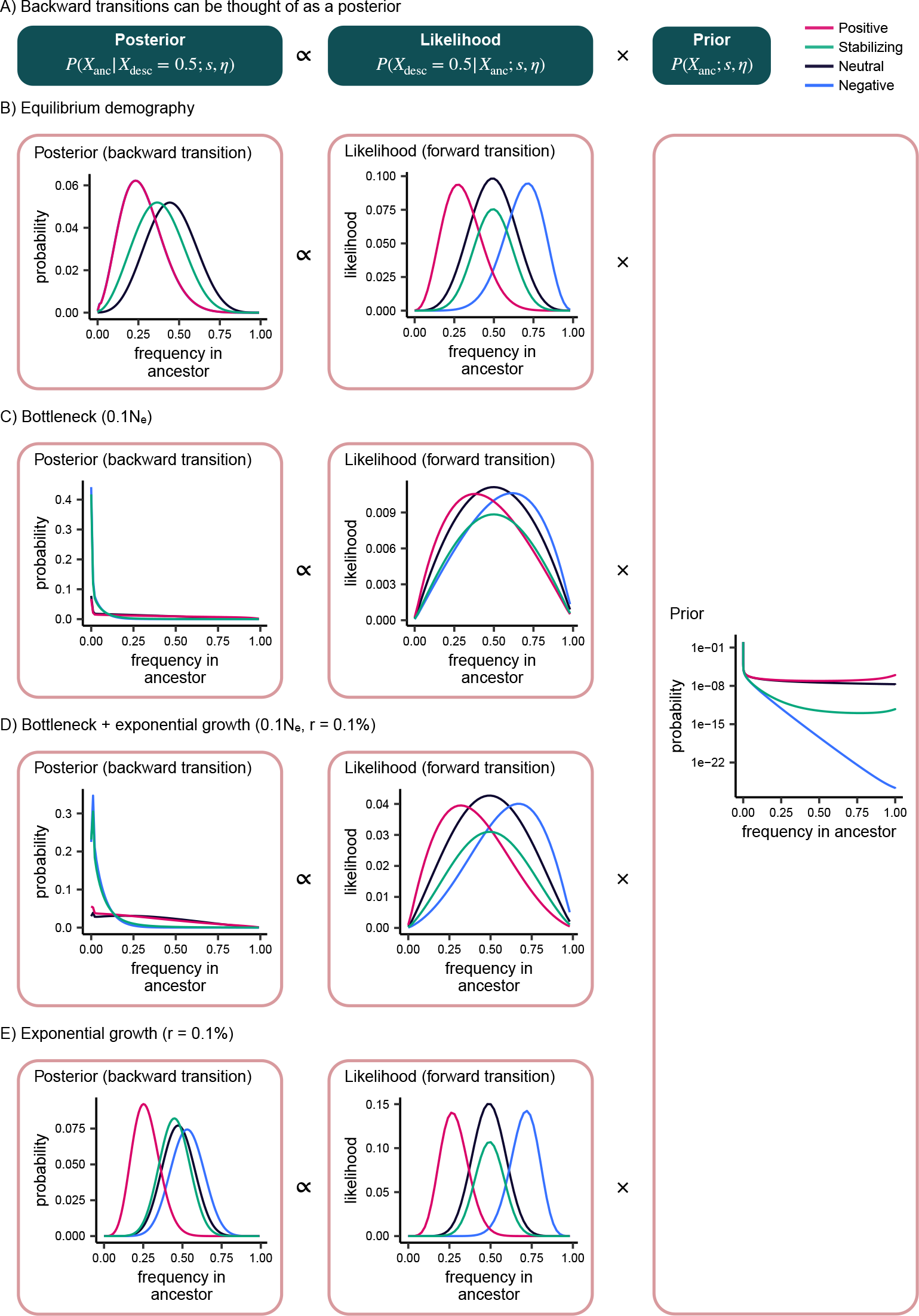
Intuition for backward transitions under different demographic scenarios. **A)** Overview of backward transition probabilities. **B-E)** Posterior, likelihood, and prior distributions for **B)** constant population size for 2,000 generations; **C)** 0.1*N_e_* bottleneck; **D)** 0.1*N_e_* bottleneck and exponential growth at a rate of 0.1%; **E)** exponential growth at a rate of 0.1%.

**Figure S3:**
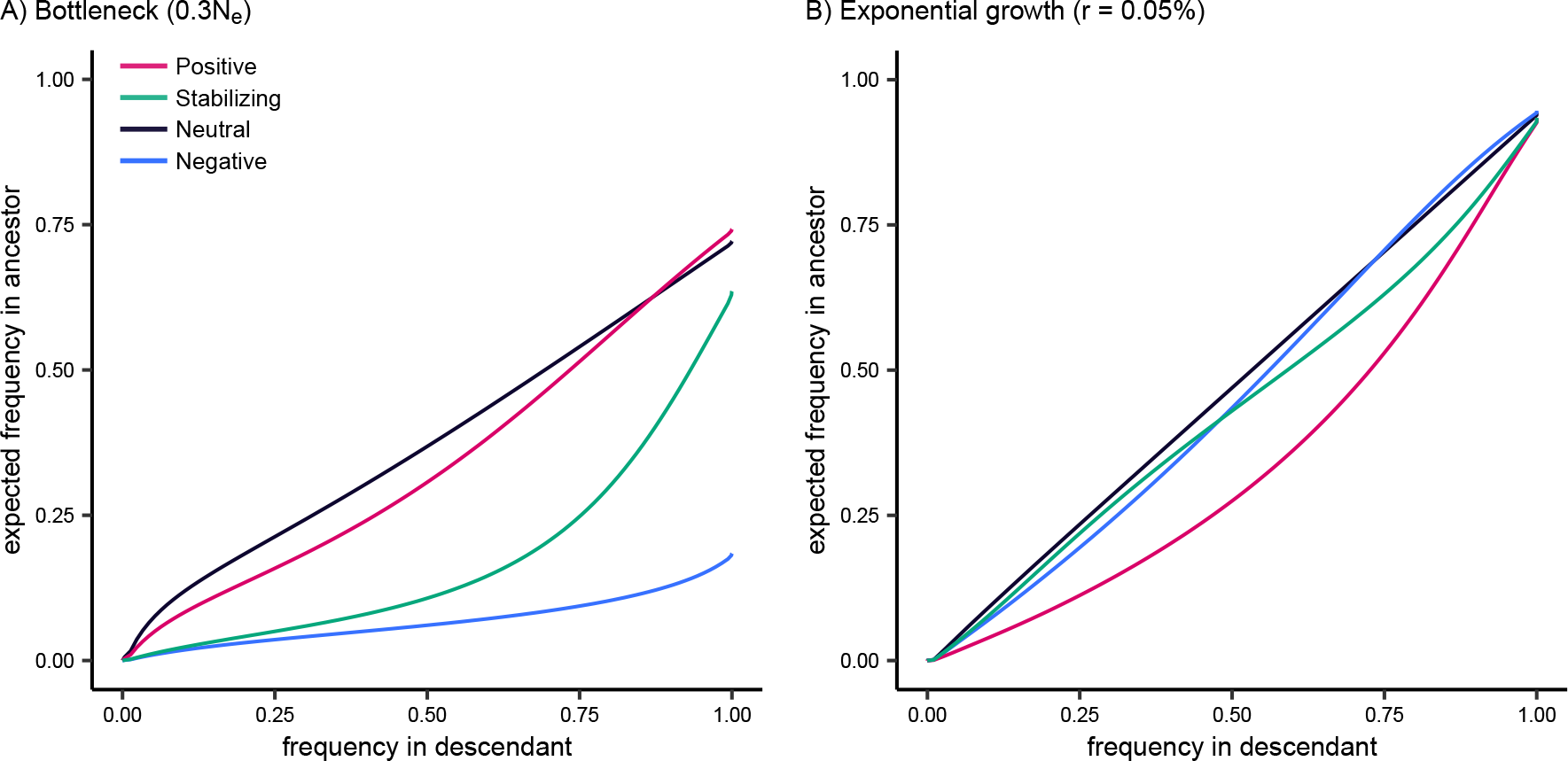
Backward transitions under additional demographic models. Expected frequency in an ancestral population conditional on the frequency in the descendant population, computed with fastDTWF. Selection coefficients correspond to |*hs*| = 5.0 × 10*^−^*^4^. Demographic models consist of an ancestral population with *N_e_* = 10,000 and **A)** 0.3*N_e_* bottleneck; **B)** exponential growth at a rate of 0.5 % each generation.

**Figure S4:**
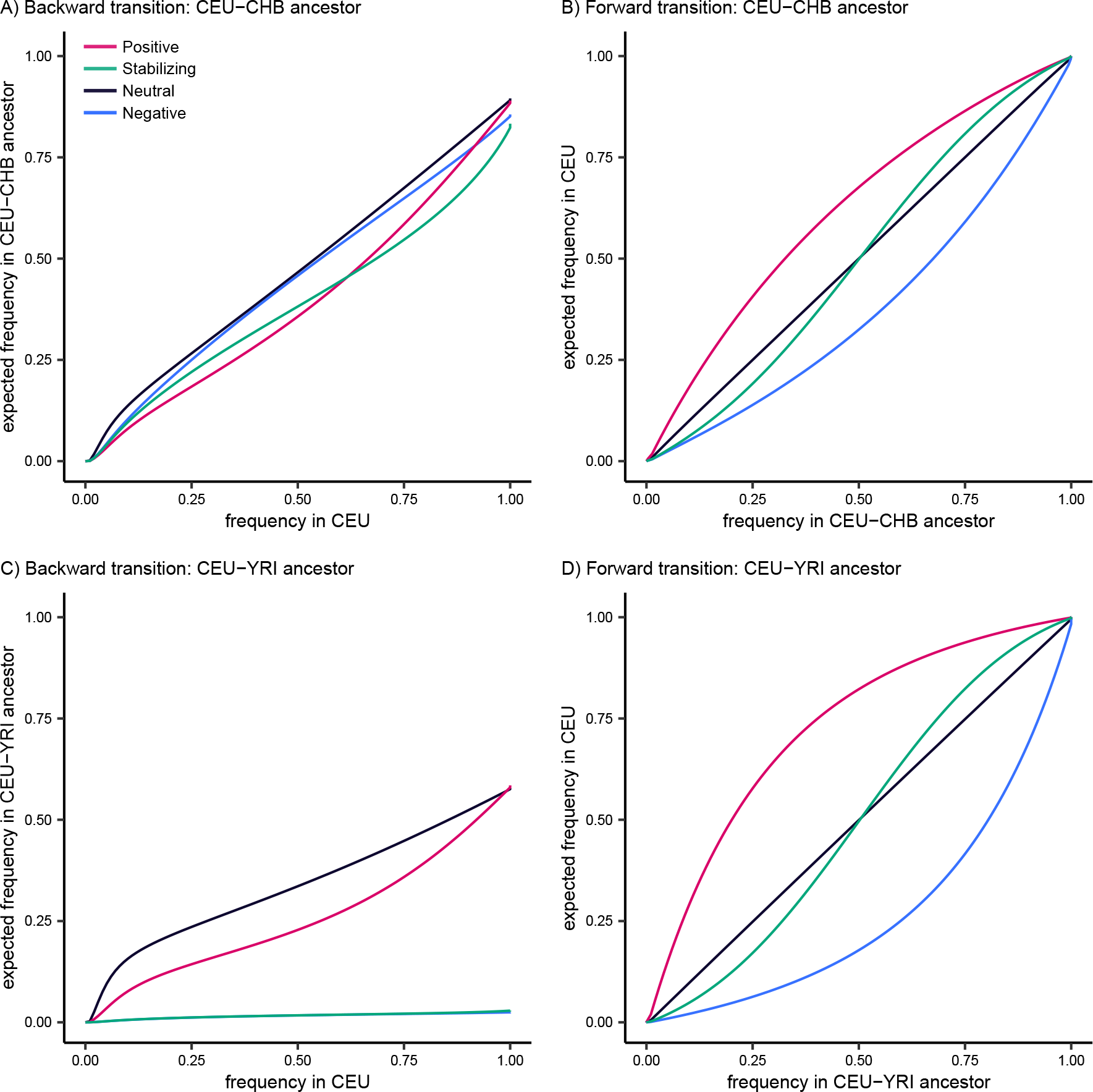
Forward and backward transitions under the out-of-Africa demographic model. **A)** Expected frequency in the CEU-CHB ancestor conditional on the frequency in CEU. **B)** Expected frequency in CEU conditional on the frequency in the CEU-CHB ancestor. **C)** Expected frequency in the CEU-YRI ancestor conditional on the frequency in CEU. **D)** Expected frequency in CEU conditional on the frequency in the CEU-YRI ancestor. For all panels, selection coefficients correspond to |*hs*| = 5.0 × 10*^−^*^4^, computed with fastDTWF.

**Figure S5:**
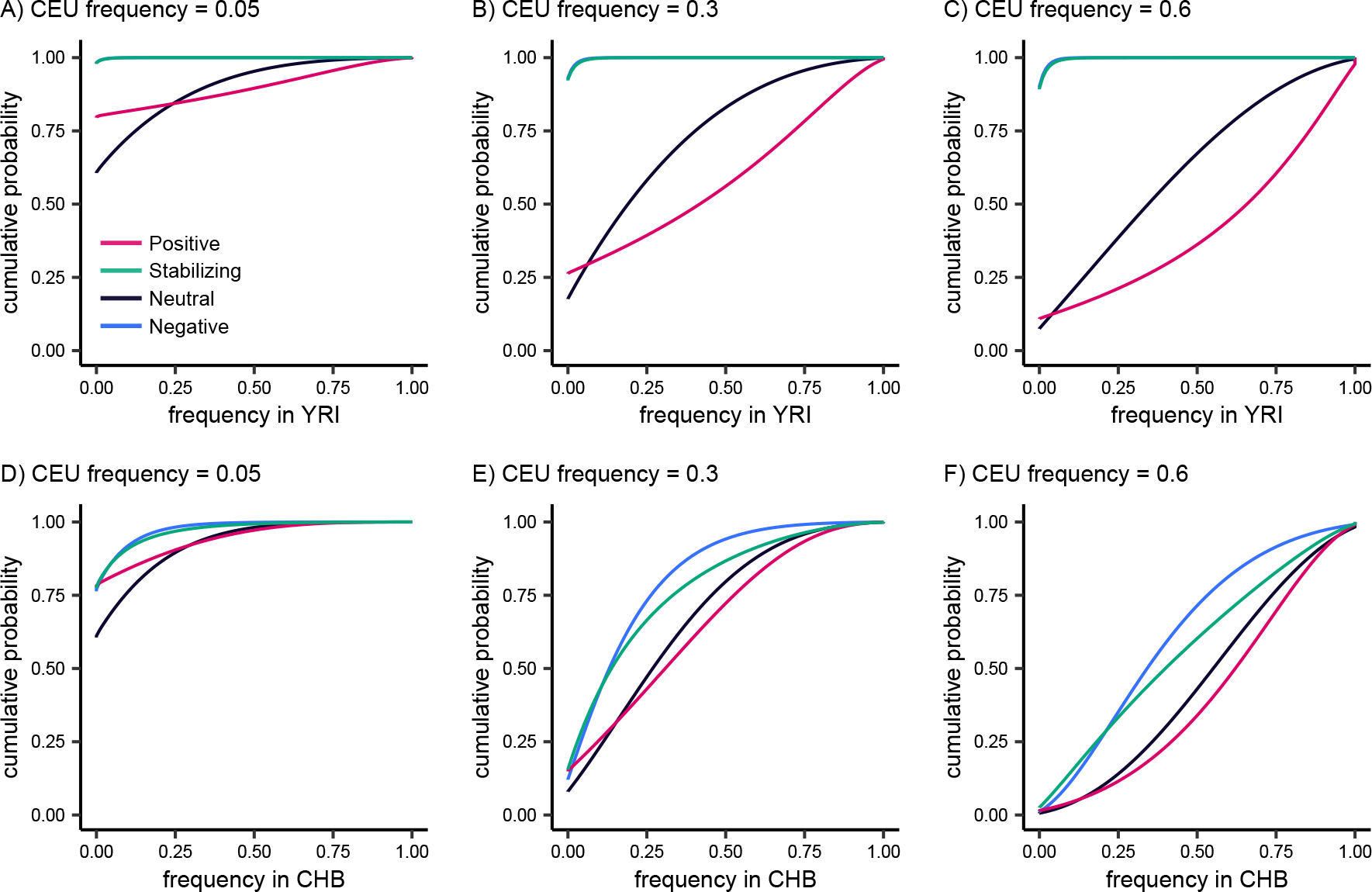
Conditional frequency spectra in YRI and CHB under strong selection. **A-C)** Cumulative probability distribution of YRI frequencies conditional on a CEU frequency of 0.05, 0.3, and 0.6. **D-F)** Cumulative probability distribution of CHB frequencies conditional on a CEU frequency of 0.05, 0.3, and 0.6. For all panels, selection coefficients correspond to *hs* = 5.0 10*^−^*^4^, computed with fastDTWF.

**Figure S6:**
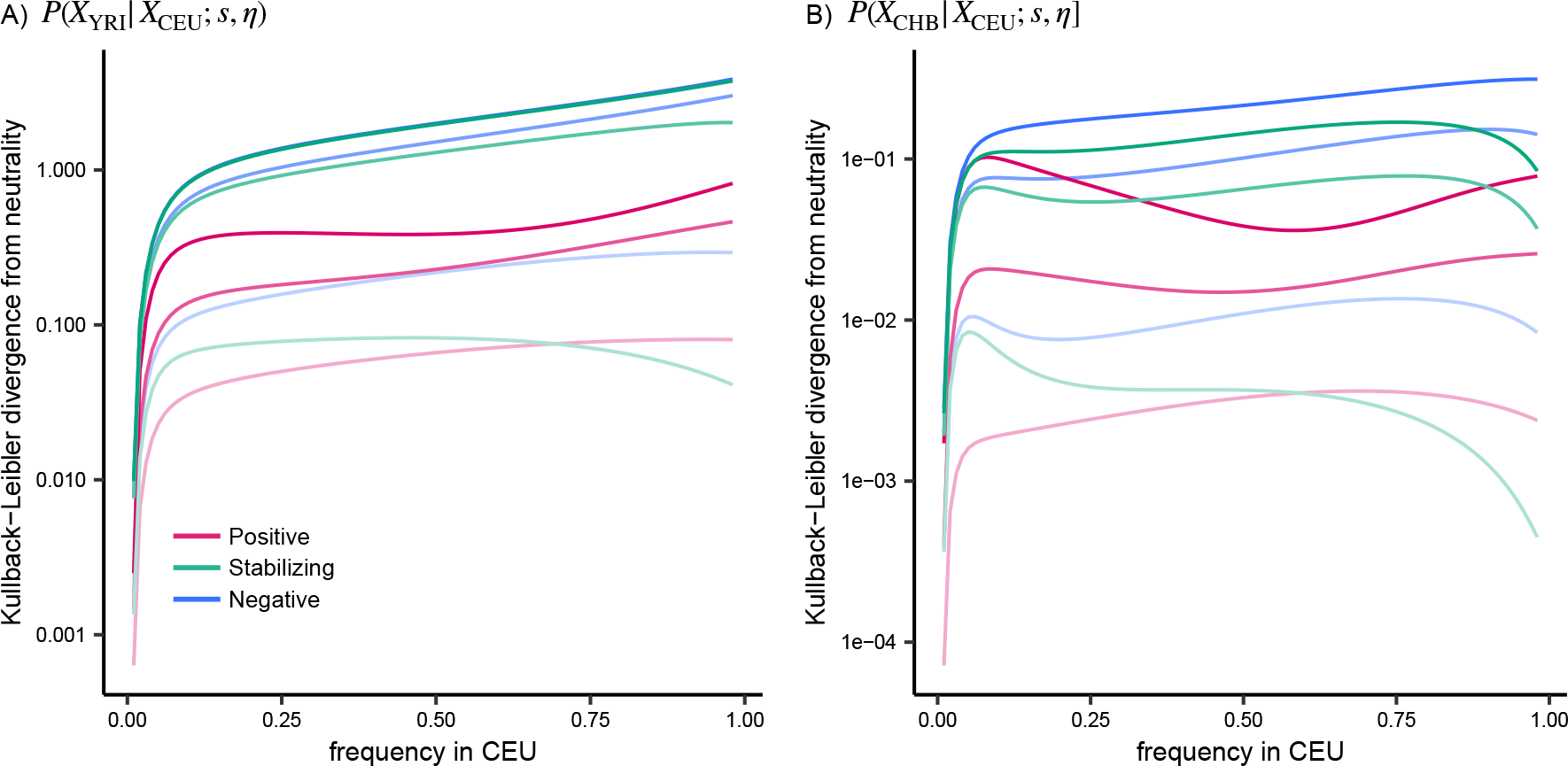
KL divergence between selection and neutrality. **A)** Kullback-Leibler divergence between the conditional frequency spectrum in YRI under neutrality and under selection, computed with fastDTWF. **B)** Kullback-Leibler divergence between the conditional frequency spectrum in CHB under neutrality and under selection, computed with fastDTWF. For both panels, selection coefficients range from *hs* = 5.0 10*^−^*^5^, shown in the lightest shades, to *hs* = 5.0 10*^−^*^4^, shown in the darkest shades.

**Figure S7:**
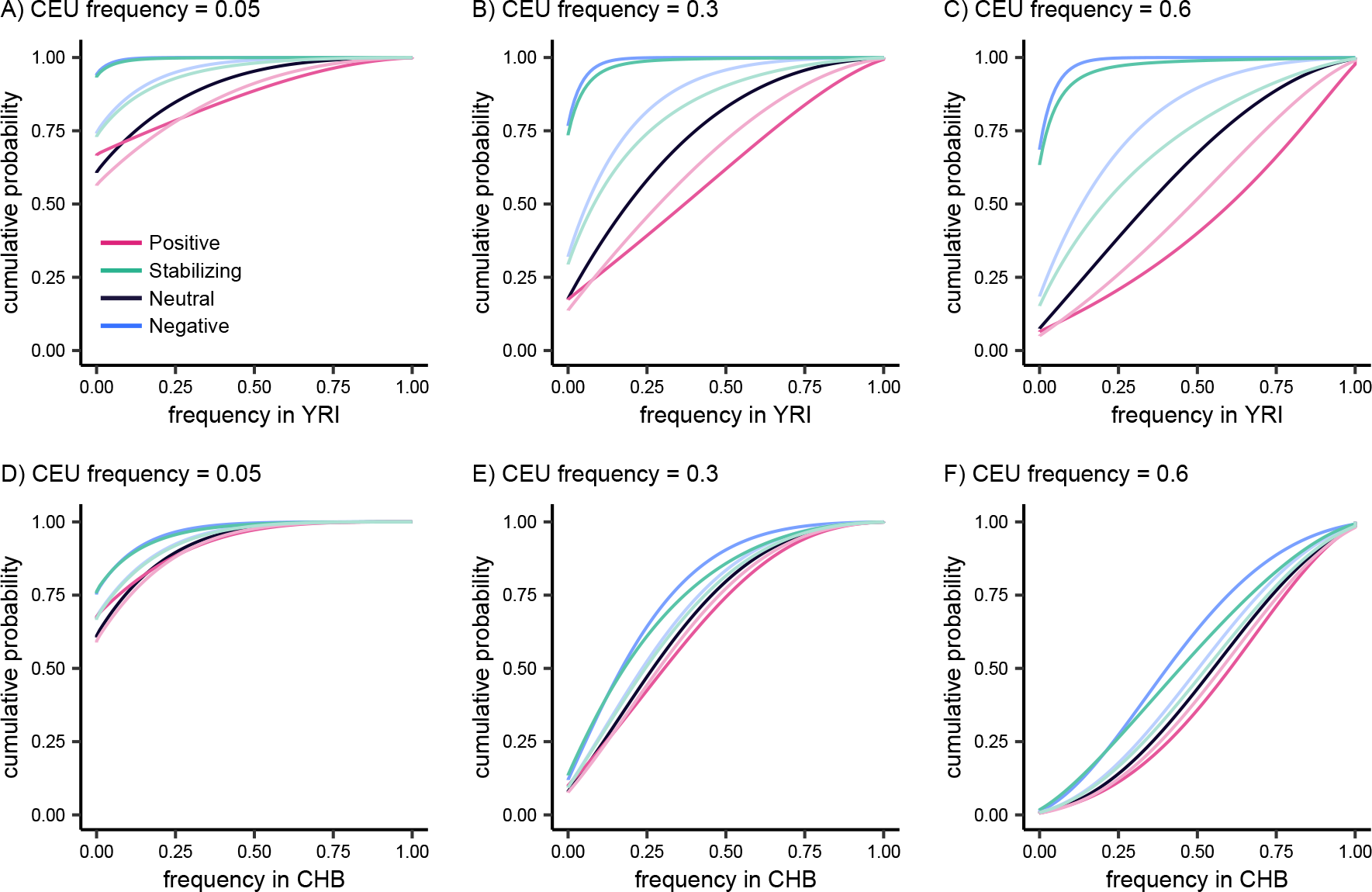
Conditional frequency spectra in YRI and CHB under weaker selection. **A-C)** Cumulative probability distribution of YRI frequencies conditional on a CEU frequency of 0.05, 0.3, and 0.6. **D-F)** Cumulative probability distribution of CHB frequencies conditional on a CEU frequency of 0.05, 0.3, and 0.6. For all panels, selection coefficients range from |*hs*| = 5.0 × 10*^−^*^5^, shown in the lightest shades, to |*hs*| = 2.5 × 10*^−^*^4^, shown in the darkest shades.

**Figure S8:**
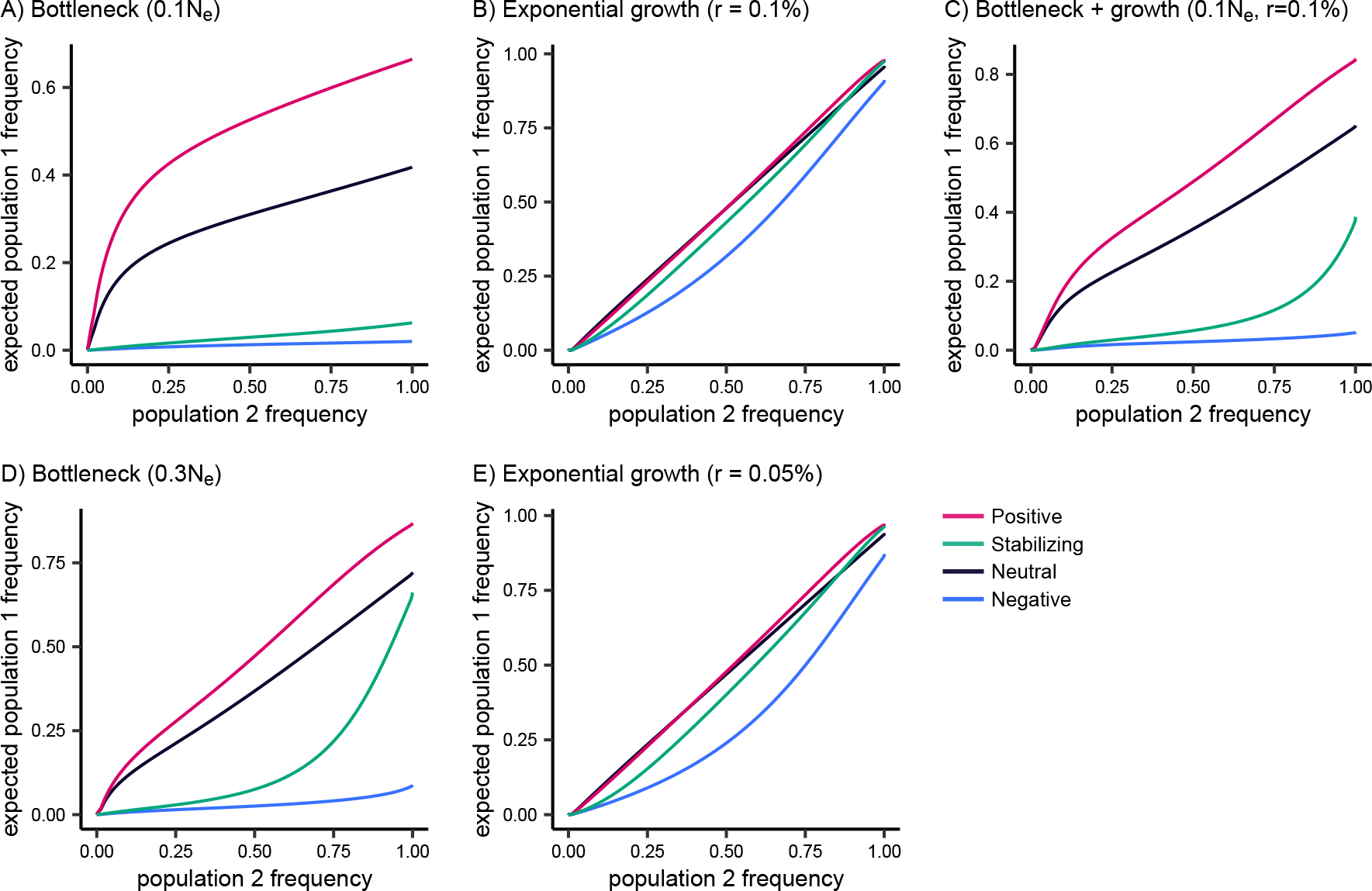
Conditional frequency spectra under different demographic scenarios. Ex-pected frequency in one descendant population conditional on another. For all panels, selection coefficients correspond to |*hs*| = 5.0 × 10*^−^*^4^, computed with fastDTWF. Demographic models consist of two populations that split from an ancestral population with *N_e_* = 10,000, 2,000 generations ago. Population 1 experiences constant population size, and population 2 (i.e. the conditional population) experiences **A)** 0.1*N_e_* bottleneck; **B)** exponential growth at a rate of 0.1% each generation; **C)** 0.1*N_e_* bottleneck and exponential growth at a rate of 0.1%; **D)** 0.3*N_e_* bottleneck; **E)** exponential growth at a rate of 0.05% each generation.

**Figure S9:**
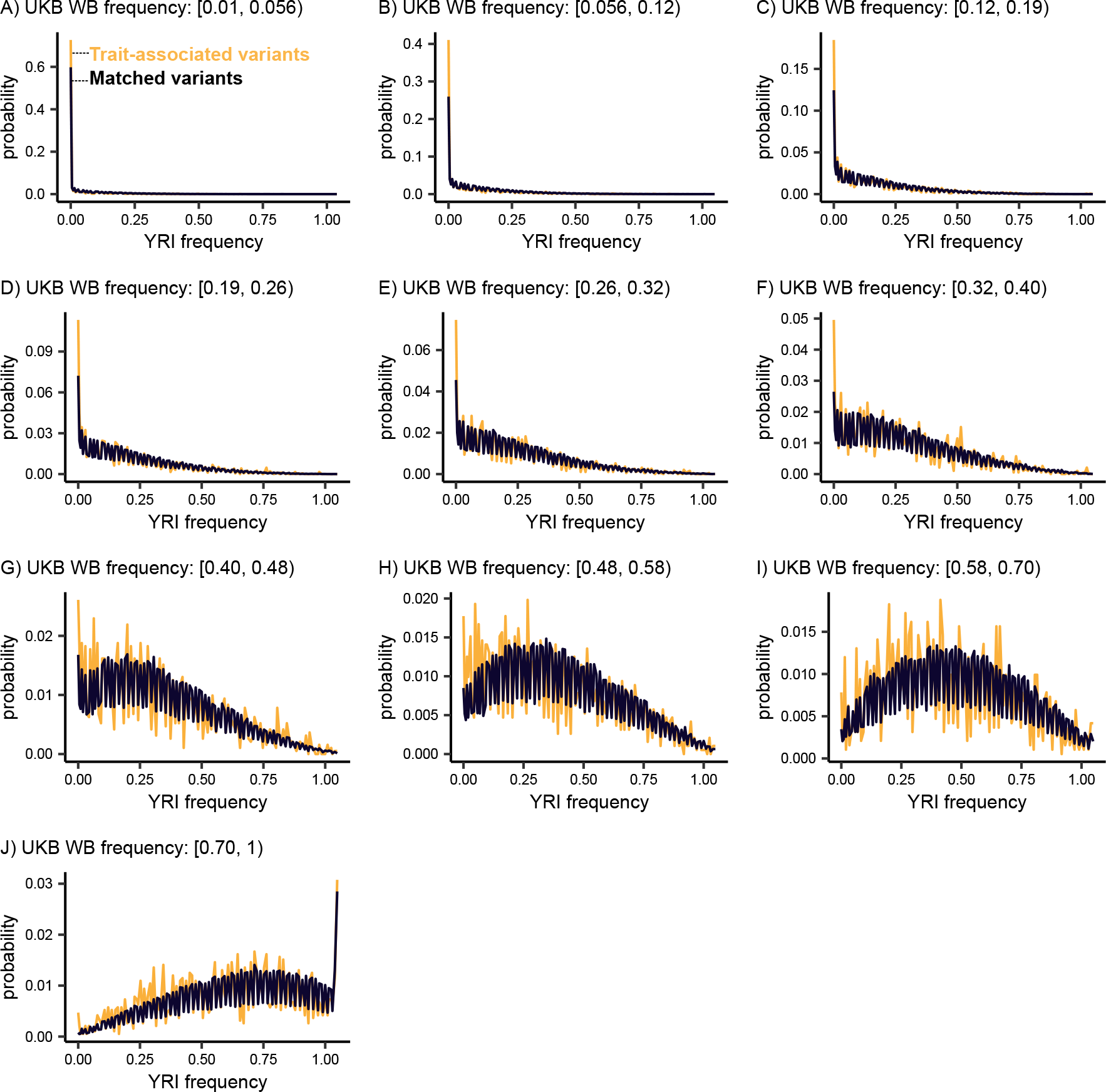
Empirical conditional frequency spectra in YRI. Distribution of YRI frequencies for all trait-associated variants and matched variants across deciles of UK Biobank White British frequency.

**Figure S10:**
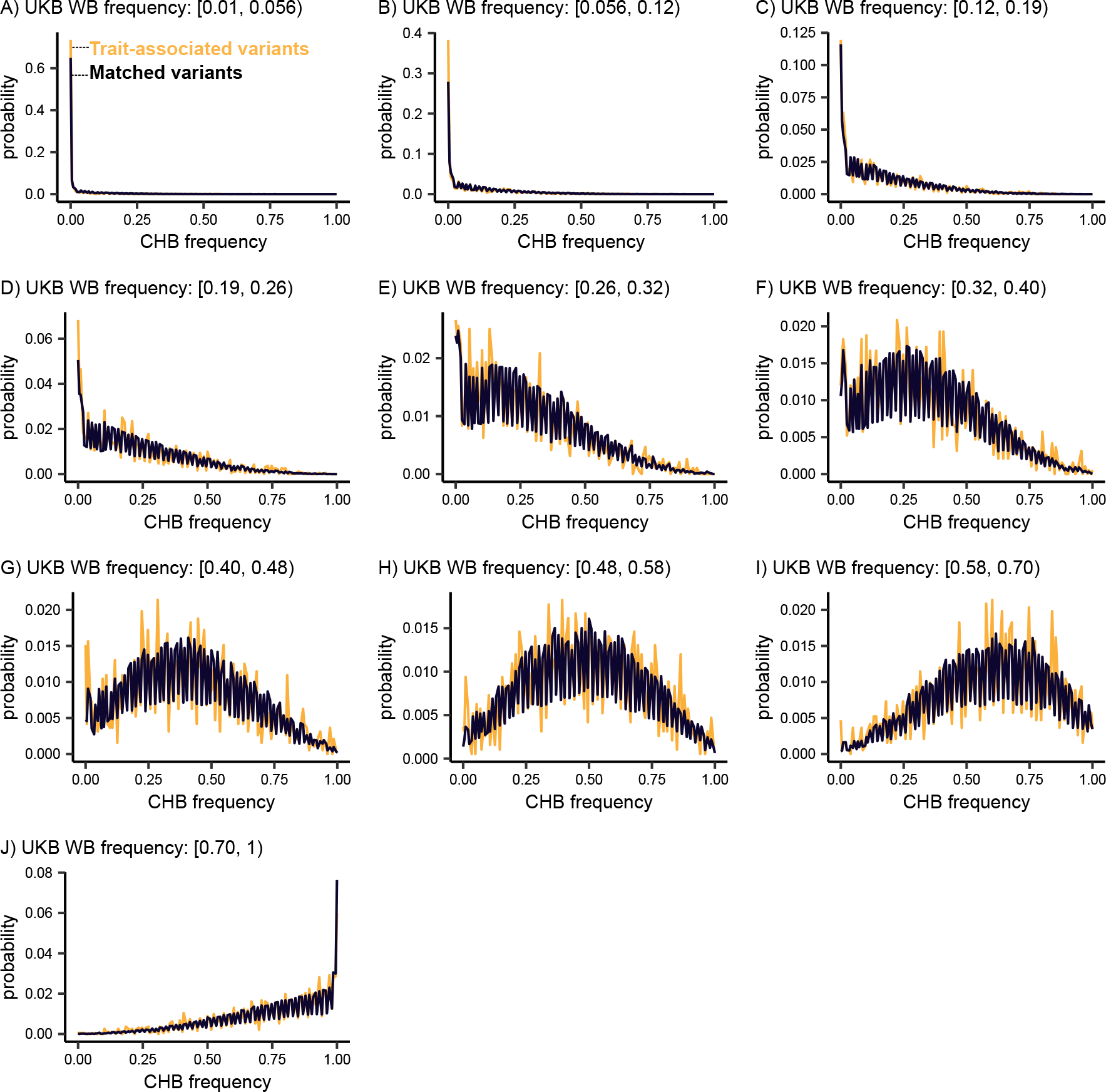
Empirical conditional frequency spectra in CHB. Distribution of CHB frequencies for all trait-associated variants and matched variants across deciles of UK Biobank White British frequency.

**Figure S11:**
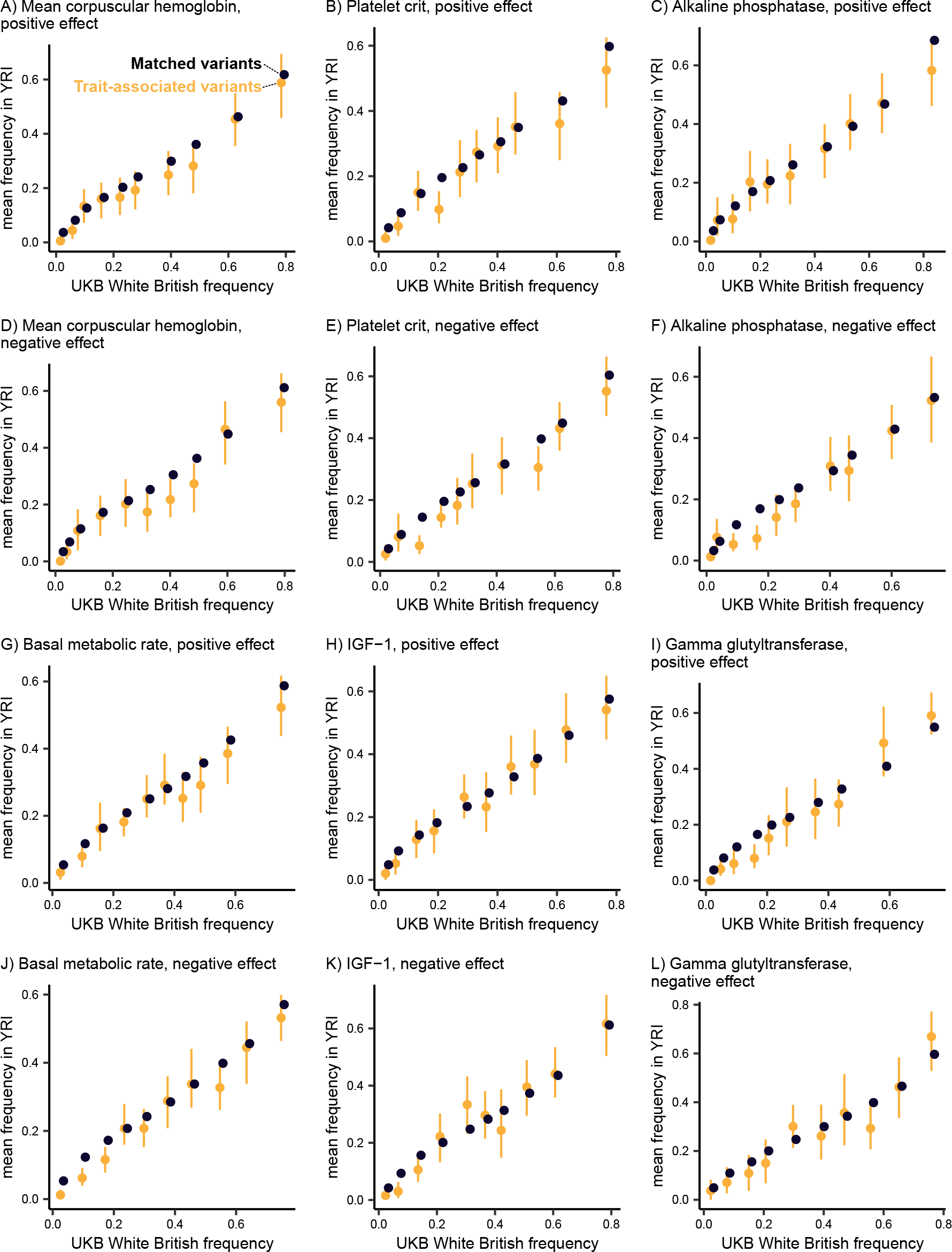
Empirical conditional frequency spectra for additional traits. Mean frequency in YRI conditional on UK Biobank White British frequency decile for variants associated with mean corpuscular hemoglobin, platelet crit, alkaline phosphatase, basal metabolic rate, IGF-1, and gamma glutyltransferase. Error bars depict the 95% confidence interval for the mean, calculated from 100 bootstrap samples. Points are jittered along the x-axis (UK Biobank White British frequency) for better visibility.

**Figure S12:**
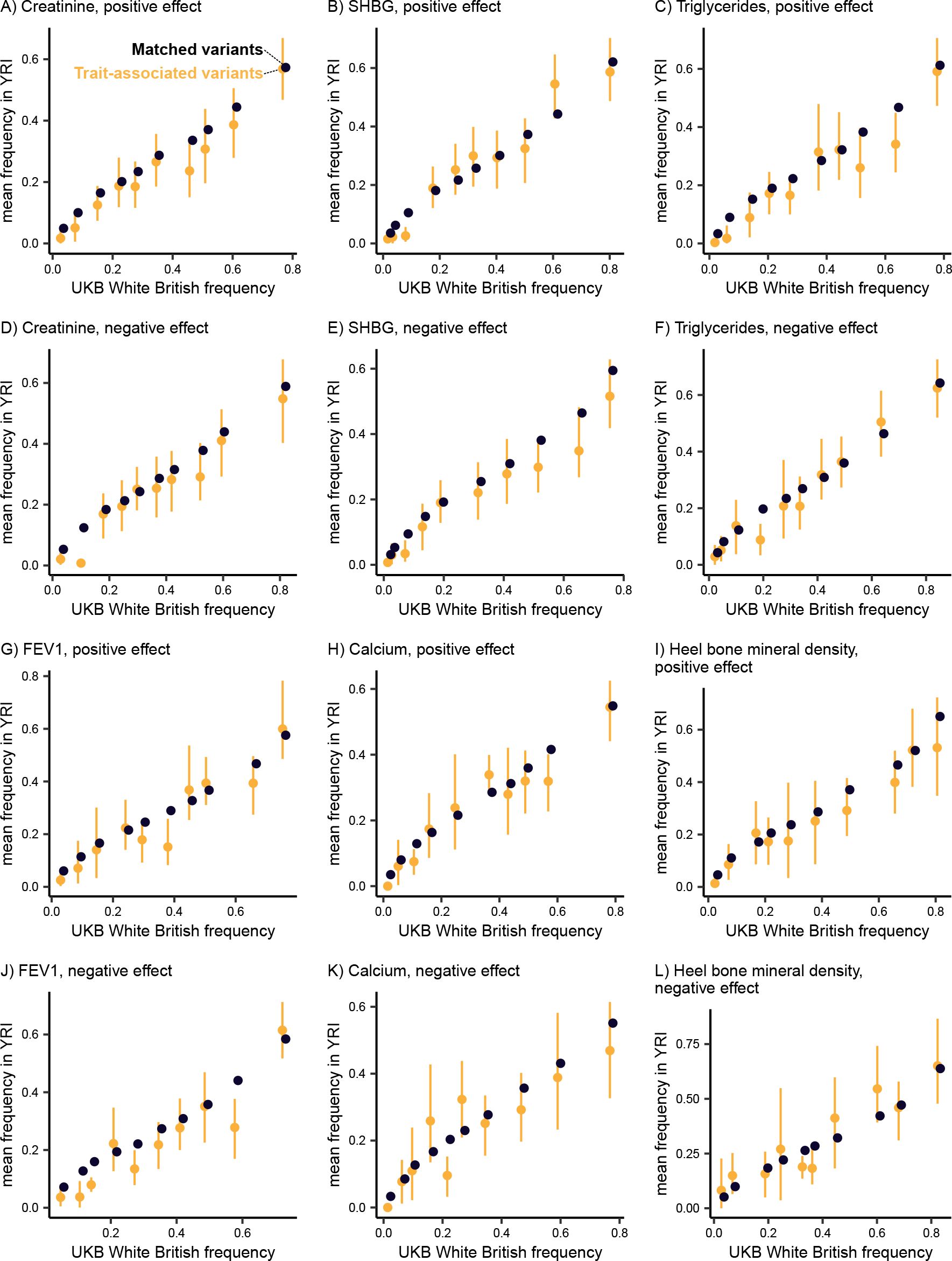
Empirical conditional frequency spectra for additional traits. Mean frequency in YRI conditional on UK Biobank White British frequency decile for variants associated with cre-atinine, SHBG, triglycerides, FEV1, calcium, and heel bone mineral density. Error bars depict the 95% confidence interval for the mean, calculated from 100 bootstrap samples. Points are jittered along the x-axis (UK Biobank White British frequency) for better visibility.

**Figure S13:**
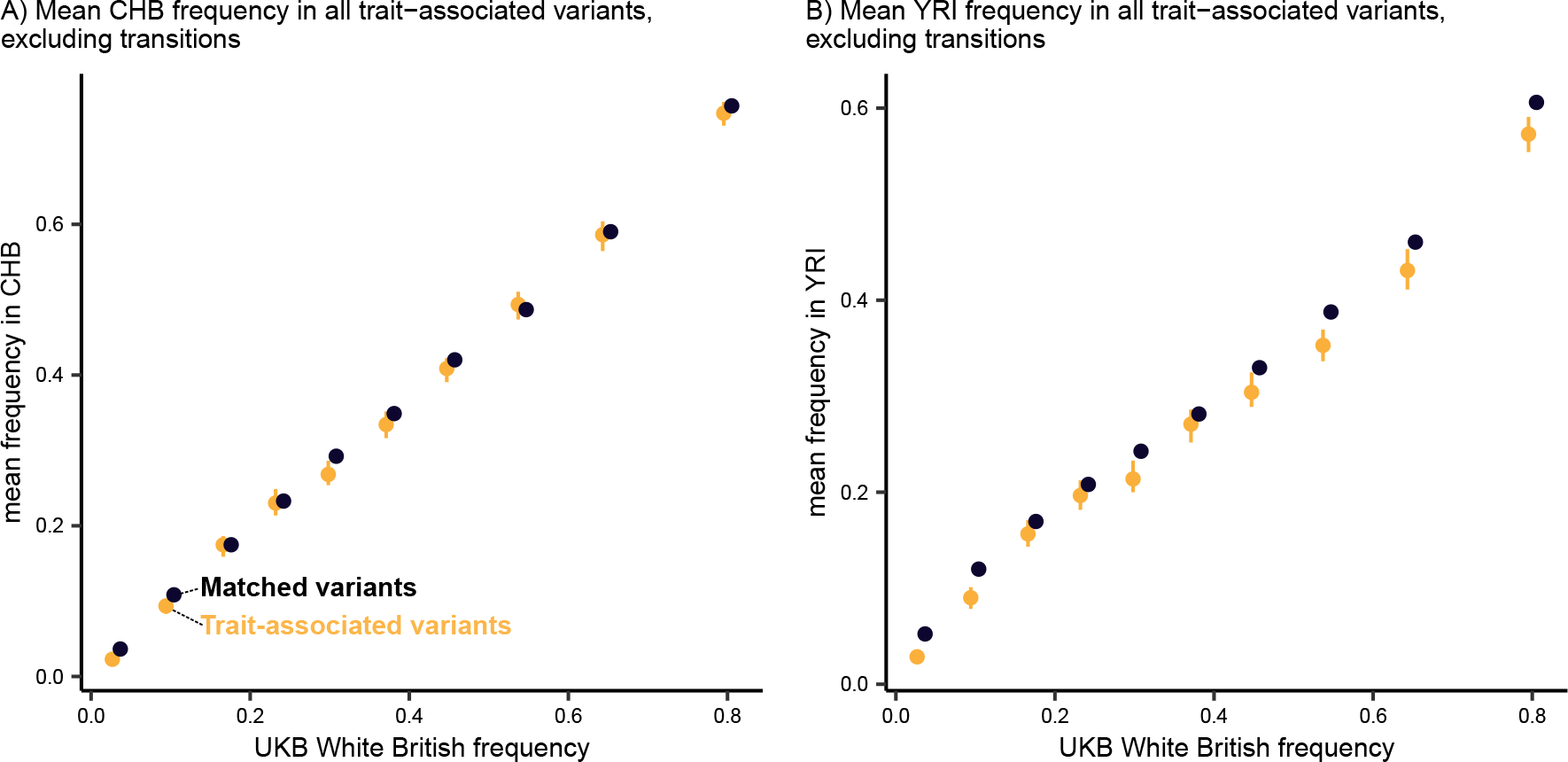
Empirical conditional frequency spectra, excluding transitions. Mean frequency in **A)** CHB and **B)** YRI conditional on UK Biobank White British frequency decile for all trait-associated variants and matched variants, excluding transition mutations. Points are jittered along the x-axis (UK Biobank White British frequency) for better visibility.

**Figure S14:**
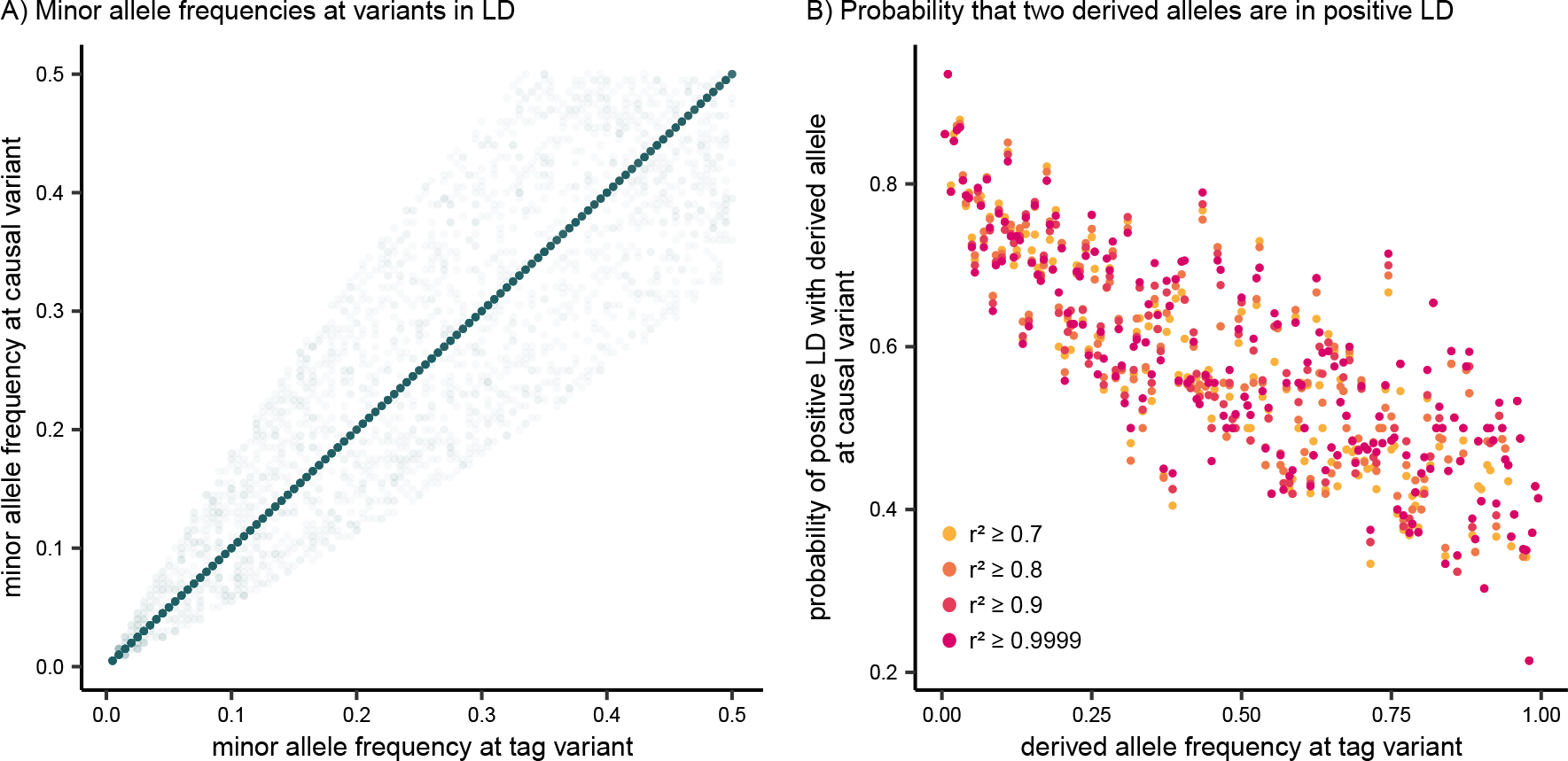
Analysis of imperfect tagging in coalescent simulations. **A)** Frequencies at pairs of variants with *r*^2^ *>* 0.5. **B)** Probability that two derived alleles are positively correlated at a pair of linked variants, conditional on the frequency at one of the variants. *r*^2^ thresholds for the linked variants range from 0.7 to 0.9999.

**Figure S15:**
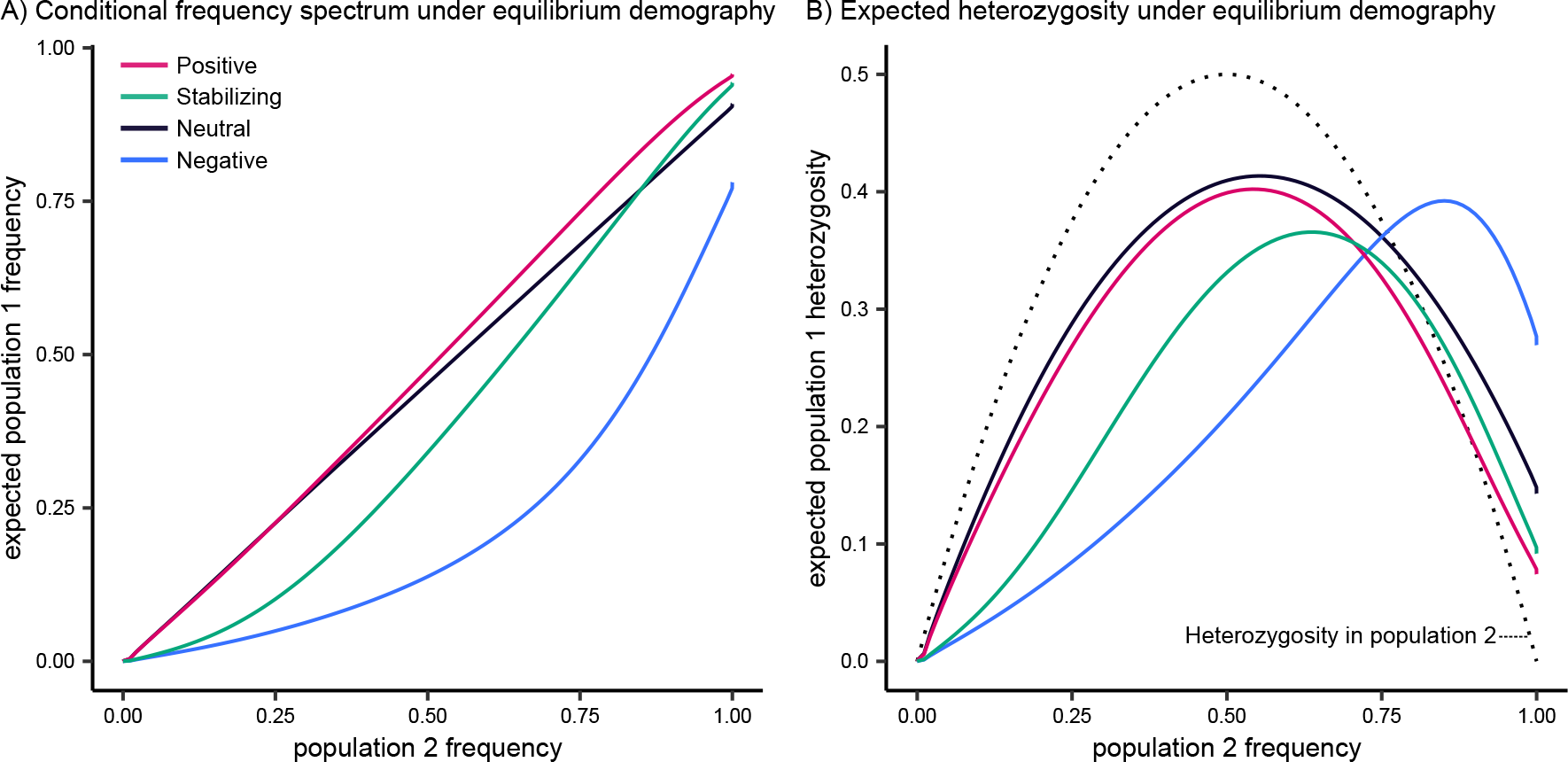
Implications for equilibrium demography. **A)** Expected frequency in one descen-dant population conditional on another. **B)** Expected heterozygosity in one descendant population conditional on another. Dotted line corresponds to the heterozygosity in the conditional population. For both panels, selection coefficients correspond to *hs* = 5.0 10*^−^*^4^, computed with fastDTWF. The demographic model consists of two populations that split from an ancestral population 2,000 generations ago and maintained a constant population size with *N_e_* = 10,000.

## References

Speed, D., G. Hemani, M. Johnson, and D. Balding, 2012 Improved Heritability Estimation from Genome-wide SNPs. American Journal of Human Genetics 91: 1011–1021.

Yang, J., B. Benyamin, B. P. McEvoy, S. Gordon, A. K. Henders, D. R. Nyholt, P. A. Madden, A. C. Heath, N. G. Martin, G. W. Montgomery, et al., 2010 Common SNPs explain a large proportion of the heritability for human height. Nature Genetics 42: 565–569.

Yang, J., A. Bakshi, Z. Zhu, G. Hemani, A. A. E. Vinkhuyzen, S. H. Lee, M. R. Robinson, J. R. B. Perry, I. M. Nolte, J. V. van Vliet-Ostaptchouk, et al., 2015 Genetic variance estimation with imputed variants finds negligible missing heritability for human height and body mass index. Nature Genetics 47: 1114–1120.

Schoech, A. P., D. M. Jordan, P.-R. Loh, S. Gazal, L. J. O’Connor, D. J. Balick, P. F. Palamara, H. K. Finucane, S. R. Sunyaev, and A. L. Price, 2019 Quantification of frequency-dependent genetic architectures in 25 UK Biobank traits reveals action of negative selection. Nature Communications 10: 790.

Zeng, J., R. de Vlaming, Y. Wu, M. R. Robinson, L. R. Lloyd-Jones, L. Yengo, C. X. Yap, A. Xue, J. Sidorenko, A. F. McRae, et al., 2018 Signatures of negative selection in the genetic architecture of human complex traits. Nature Genetics 50: 746–753.

Zeng, J., A. Xue, L. Jiang, L. R. Lloyd-Jones, Y. Wu, H. Wang, Z. Zheng, L. Yengo, K. E. Kemper, M. E. Goddard, et al., 2021 Widespread signatures of natural selection across human complex traits and functional genomic categories. Nature Communications 12: 1–12.

Gazal, S., H. K. Finucane, N. A. Furlotte, P.-R. Loh, P. F. Palamara, X. Liu, A. Schoech, B. Bulik-Sullivan, B. M. Neale, A. Gusev, et al., 2017 Linkage disequilibrium-dependent architecture of human complex traits shows action of negative selection. Nature Genetics 49: 1421–1427.

Speed, D., J. Holmes, and D. J. Balding, 2020 Evaluating and improving heritability models using summary statistics. Nature Genetics 52: 458–462.

Lande, R., 1976 Natural Selection and Random Genetic Drift in Phenotypic Evolution. Evolution 30: 314–334.

Simons, Y. B., K. Bullaughey, R. R. Hudson, and G. Sella, 2018 A population genetic interpretation of GWAS findings for human quantitative traits. PLoS Biology 16: e2002985.

Turelli, M., 1984 Heritable genetic variation via mutation-selection balance: Lerch’s zeta meets the abdominal bristle. Theoretical Population Biology 25: 138–193.

Robertson, A., 1956 The effect of selection against extreme deviants based on deviation or on homozygosis. Journal of Genetics 54: 236–248.

Walsh, B. and M. Lynch, 2018 Evolution and Selection of Quantitative Traits. Oxford University Press.

Caballero, A., A. Tenesa, and P. D. Keightley, 2015 The Nature of Genetic Variation for Complex Traits Revealed by GWAS and Regional Heritability Mapping Analyses. Genetics 201: 1601–1613.

Eyre-Walker, A., 2010 Genetic architecture of a complex trait and its implications for fitness and genome-wide association studies. Proceedings of the National Academy of Sciences 107: 1752–1756.

Keightley, P. D. and W. G. Hill, 1990 Variation Maintained in Quantitative Traits with Mutation-Selection Balance: Pleiotropic Side-Effects on Fitness Traits. Proceedings: Biological Sciences 242: 95–100.

Guo, J., Y. Wu, Z. Zhu, Z. Zheng, M. Trzaskowski, J. Zeng, M. R. Robinson, P. M. Visscher, and J. Yang, 2018 Global genetic differentiation of complex traits shaped by natural selection in humans. Nature Communications 9: 1865.

Mostafavi, H., J. P. Spence, S. Naqvi, and J. K. Pritchard, 2023 Systematic differences in discovery of genetic effects on gene expression and complex traits. Nature Genetics 55: 1866–1875.

O’Connor, L. J., A. P. Schoech, F. Hormozdiari, S. Gazal, N. Patterson, and A. L. Price, 2019 Extreme polygenicity of complex traits is explained by negative selection. American Journal of Human Genetics 105: 456–476.

Weiner, D. J., A. Nadig, K. A. Jagadeesh, K. K. Dey, B. M. Neale, E. B. Robinson, K. J. Karczewski, and L. J. O’Connor, 2023 Polygenic architecture of rare coding variation across 394,783 exomes. Nature 614: 492–499.

Durvasula, A. and K. E. Lohmueller, 2021 Negative selection on complex traits limits phenotype prediction accuracy between populations. American Journal of Human Genetics 108: 620–631.

Wang, Y., J. Guo, G. Ni, J. Yang, P. M. Visscher, and L. Yengo, 2020 Theoretical and empirical quantification of the accuracy of polygenic scores in ancestry divergent populations. Nature Communications 11: 3865.

Yair, S. and G. Coop, 2022 Population differentiation of polygenic score predictions under stabilizing selection. Philosophical Transactions of the Royal Society B: Biological Sciences 377: 20200416.

Manolio, T. A., F. S. Collins, N. J. Cox, D. B. Goldstein, L. A. Hindorff, D. J. Hunter, M. I. McCarthy, E. M. Ramos, L. R. Cardon, A. Chakravarti, et al., 2009 Finding the missing heritability of complex diseases. Nature 461: 747–753.

Pritchard, J. K., 2001 Are rare variants responsible for susceptibility to complex diseases? American Journal of Human Genetics 69: 124–137.

Clark, A. G., M. J. Hubisz, C. D. Bustamante, S. H. Williamson, and R. Nielsen, 2005 Ascertainment bias in studies of human genome-wide polymorphism. Genome Research 15: 1496–1502.

Lachance, J. and S. A. Tishkoff, 2013 SNP ascertainment bias in population genetic analyses: Why it is important, and how to correct it. BioEssays 35: 780–786.

Maruyama, T., 1974 The age of an allele in a finite population. Genetical Research 23: 137–143.

Baharian, S. and S. Gravel, 2018 On the decidability of population size histories from finite allele frequency spectra. Theoretical Population Biology 120: 42–51.

Bhaskar, A. and Y. S. Song, 2014 Descartes’ rule of signs and the identifiability of population demographic models from genomic variation data. Annals of Statistics 42: 2469–2493.

Bhaskar, A., Y. X. R. Wang, and Y. S. Song, 2015 Efficient inference of population size histories and locus-specific mutation rates from large-sample genomic variation data. Genome Research 25: 268–279.

Myers, S., C. Fefferman, and N. Patterson, 2008 Can one learn history from the allelic spectrum? Theoretical Population Biology 73: 342–348.

Ragsdale, A. P., C. Moreau, and S. Gravel, 2018 Genomic inference using diffusion models and the allele frequency spectrum. Current Opinion in Genetics & Development 53: 140–147.

Rosen, Z., A. Bhaskar, S. Roch, and Y. S. Song, 2018 Geometry of the Sample Frequency Spectrum and the Perils of Demographic Inference. Genetics 210: 665–682.

Schraiber, J. G., 2018 Assessing the Relationship of Ancient and Modern Populations. Genetics 208: 383–398.

Spence, J. P., J. A. Kamm, and Y. S. Song, 2016 The Site Frequency Spectrum for General Coalescents. Genetics 202: 1549–1561.

Terhorst, J. and Y. S. Song, 2015 Fundamental limits on the accuracy of demographic inference based on the sample frequency spectrum. Proceedings of the National Academy of Sciences 112: 7677–7682.

Chen, H. and M. Slatkin, 2013 Inferring Selection Intensity and Allele Age from Multilocus Haplotype Structure. G3: Genes, Genomes, Genetics 3: 1429–1442.

Evans, S. N., Y. Shvets, and M. Slatkin, 2007 Non-equilibrium theory of the allele frequency spectrum. Theoretical Population Biology 71: 109–119.

Schraiber, J. G., S. N. Evans, and M. Slatkin, 2016 Bayesian Inference of Natural Selection from Allele Frequency Time Series. Genetics 203: 493–511.

Slatkin, M., 2001 Simulating genealogies of selected alleles in a population of variable size. Genetical Research 78: 49–57.

Song, Y. S. and M. Steinrücken, 2012 A Simple Method for Finding Explicit Analytic Transition Densities of Diffusion Processes with General Diploid Selection. Genetics 190: 1117–1129.

΁ivković, D. and W. Stephan, 2011 Analytical results on the neutral non-equilibrium allele frequency spectrum based on diffusion theory. Theoretical Population Biology 79: 184–191.

Dilber, E. and J. Terhorst, 2024 Faster inference of complex demographic models from large allele frequency spectra.

Gutenkunst, R. N., R. D. Hernandez, S. H. Williamson, and C. D. Bustamante, 2009 Inferring the Joint Demographic History of Multiple Populations from Multidimensional SNP Frequency Data. PLoS Genetics 5: e1000695.

Jouganous, J., W. Long, A. P. Ragsdale, and S. Gravel, 2017 Inferring the Joint Demographic History of Multiple Populations: Beyond the Diffusion Approximation. Genetics 206: 1549–1567.

Kamm, J. A., J. Terhorst, and Y. S. Song, 2017 Efficient Computation of the Joint Sample Frequency Spectra for Multiple Populations. Journal of Computational and Graphical Statistics 26: 182–194.

Kamm, J., J. Terhorst, R. Durbin, and Y. S. Song, 2020 Efficiently Inferring the Demographic History of Many Populations With Allele Count Data. Journal of the American Statistical Association 115: 1472–1487.

Kern, A. D. and J. Hey, 2017 Exact Calculation of the Joint Allele Frequency Spectrum for Isolation with Migration Models. Genetics 207: 241–253.

Lukić, S., J. Hey, and K. Chen, 2011 Non-equilibrium allele frequency spectra via spectral methods. Theoretical Population Biology 79: 203–219.

Lukić, S. and J. Hey, 2012 Demographic Inference Using Spectral Methods on SNP Data, with an Analysis of the Human Out-of-Africa Expansion. Genetics 192: 619–639.

Yang, M. A., K. Harris, and M. Slatkin, 2014 The Projection of a Test Genome onto a Reference Population and Applications to Humans and Archaic Hominins. Genetics 198: 1655–1670.

Harpak, A., A. Bhaskar, and J. K. Pritchard, 2016 Mutation Rate Variation is a Primary Determinant of the Distribution of Allele Frequencies in Humans. PLoS Genetics 12: e1006489.

Durvasula, A. and S. Sankararaman, 2020 Recovering signals of ghost archaic introgression in African populations. Science Advances 6: eaax5097.

Haller, B. C. and P. W. Messer, 2019 SLiM 3: Forward Genetic Simulations Beyond the Wright-Fisher Model. Molecular Biology and Evolution 36: 632–637.

Spence, J. P., T. Zeng, H. Mostafavi, and J. K. Pritchard, 2023 Scaling the discrete-time Wright–Fisher model to biobank-scale datasets. Genetics 225: iyad168.

Auton, A., G. R. Abecasis, D. M. Altshuler, R. M. Durbin, G. R. Abecasis, D. R. Bentley, A. Chakravarti, A. G. Clark, P. Donnelly, E. E. Eichler, et al., 2015 A global reference for human genetic variation. Nature 526: 68–74.

Mills, M. C. and C. Rahal, 2019 A scientometric review of genome-wide association studies. Communications Biology 2: 1–11.

Murphy, D. A., E. Elyashiv, G. Amster, and G. Sella, 2022 Broad-scale variation in human genetic diversity levels is predicted by purifying selection on coding and non-coding elements. eLife 12: e76065.

Jónsson, H., P. Sulem, B. Kehr, S. Kristmundsdottir, F. Zink, E. Hjartarson, M. T. Hardarson, K. E. Hjorleifsson, H. P. Eggertsson, S. A. Gudjonsson, et al., 2017 Parental influence on human germline de novo mutations in 1,548 trios from Iceland. Nature 549: 519–522.

Keightley, P. D. and B. C. Jackson, 2018 Inferring the Probability of the Derived vs. the Ancestral Allelic State at a Polymorphic Site. Genetics 209: 897–906.

Ding, Y., K. Hou, Z. Xu, A. Pimplaskar, E. Petter, K. Boulier, F. Privé, B. J. Vilhjálmsson, L. M. Olde Loohuis, and B. Pasaniuc, 2023 Polygenic scoring accuracy varies across the genetic ancestry continuum. Nature 618: 774–781.

Martin, A. R., M. Kanai, Y. Kamatani, Y. Okada, B. M. Neale, and M. J. Daly, 2019 Clinical use of current polygenic risk scores may exacerbate health disparities. Nature Genetics 51: 584–591.

Mostafavi, H., A. Harpak, I. Agarwal, D. Conley, J. K. Pritchard, and M. Przeworski, 2020 Variable prediction accuracy of polygenic scores within an ancestry group. eLife 9: e48376.

Patel, R. A., S. A. Musharoff, J. P. Spence, H. Pimentel, C. Tcheandjieu, H. Mostafavi, N. Sinnott-Armstrong, S. L. Clarke, C. J. Smith, V.A. Million Veteran Program, et al., 2022 Genetic interactions drive heterogeneity in causal variant effect sizes for gene expression and complex traits. American Journal of Human Genetics 109: 1286–1297.

Privé, F., H. Aschard, S. Carmi, L. Folkersen, C. Hoggart, P. F. O’Reilly, and B. J. Vilhjálmsson, 2022 Portability of 245 polygenic scores when derived from the UK Biobank and applied to 9 ancestry groups from the same cohort. The American Journal of Human Genetics 109: 12–23, Publisher: Elsevier.

Wojcik, G. L., M. Graff, K. K. Nishimura, R. Tao, J. Haessler, C. R. Gignoux, H. M. Highland, Y. M. Patel, E. P. Sorokin, C. L. Avery, et al., 2019 Genetic analyses of diverse populations improves discovery for complex traits. Nature 570: 514–518.

Vilhjálmsson, B. J., J. Yang, H. K. Finucane, A. Gusev, S. Lindström, S. Ripke, G. Genovese, P.-R. Loh, G. Bhatia, R. Do, et al., 2015 Modeling linkage disequilibrium increases accuracy of polygenic risk scores. American Journal of Human Genetics 97: 576–592.

Ragsdale, A. P. and S. Gravel, 2019 Models of archaic admixture and recent history from two-locus statistics. PLoS Genetics 15: e1008204.

Gillespie, J. H., 2004 Population Genetics. Johns Hopkins University Press.

Simons, Y. B., H. Mostafavi, C. J. Smith, J. K. Pritchard, and G. Sella, 2022 Simple scaling laws control the genetic architectures of human complex traits.

Aragam, K. G., T. Jiang, A. Goel, S. Kanoni, B. N. Wolford, D. S. Atri, E. M. Weeks, M. Wang, G. Hindy, W. Zhou, et al., 2022 Discovery and systematic characterization of risk variants and genes for coronary artery disease in over a million participants. Nature Genetics 54: 1803–1815.

Bellenguez, C., F. Küçükali, I. E. Jansen, L. Kleineidam, S. Moreno-Grau, N. Amin, A. C. Naj, R. Campos-Martin, B. Grenier-Boley, V. Andrade, et al., 2022 New insights into the genetic etiology of Alzheimer’s disease and related dementias. Nature Genetics 54: 412–436.

International IBD Genetics Consortium (IIBDGC), C. Agliardi, L. Alfredsson, M. Alizadeh, C. Anderson, R. Andrews, H. B. Søndergaard, A. Baker, G. Band, and others, 2013 Analysis of immune-related loci identifies 48 new susceptibility variants for multiple sclerosis. Nature Genetics 45: 1353–1360.

De Lange, K. M., L. Moutsianas, J. C. Lee, C. A. Lamb, Y. Luo, N. A. Kennedy, L. Jostins, D. L. Rice, J. Gutierrez-Achury, S.-G. Ji, et al., 2017 Genome-wide association study implicates immune activation of multiple integrin genes in inflammatory bowel disease. Nature Genetics 49: 256–261.

Demontis, D., G. B. Walters, G. Athanasiadis, R. Walters, K. Therrien, T. T. Nielsen, L. Farajzadeh, G. Voloudakis, J. Bendl, B. Zeng, et al., 2023 Genome-wide analyses of ADHD identify 27 risk loci, refine the genetic architecture and implicate several cognitive domains. Nature Genetics 55: 198–208.

Ishigaki, K., S. Sakaue, C. Terao, Y. Luo, K. Sonehara, K. Yamaguchi, T. Amariuta, C. L. Too, V. A. Laufer, I. C. Scott, et al., 2022 Multi-ancestry genome-wide association analyses identify novel genetic mechanisms in rheumatoid arthritis. Nature Genetics 54: 1640–1651.

Michailidou, K., J. Beesley, S. Lindstrom, S. Canisius, J. Dennis, M. J. Lush, M. J. Maranian, M. K. Bolla, Q. Wang, M. Shah, et al., 2015 Genome-wide association analysis of more than 120,000 individuals identifies 15 new susceptibility loci for breast cancer. Nature Genetics 47: 373–380.

Mullins, N., A. J. Forstner, K. S. O’Connell, B. Coombes, J. R. Coleman, Z. Qiao, T. D. Als, T. B. Bigdeli, S. Børte, J. Bryois, et al., 2021 Genome-wide association study of more than 40,000 bipolar disorder cases provides new insights into the underlying biology. Nature Genetics 53: 817–829.

Nalls, M. A., C. Blauwendraat, C. L. Vallerga, K. Heilbron, S. Bandres-Ciga, D. Chang, M. Tan, D. A. Kia, A. J. Noyce, A. Xue, et al., 2019 Identification of novel risk loci, causal insights, and heritable risk for Parkinson’s disease: a meta-analysis of genome-wide association studies. The Lancet Neurology 18: 1091–1102.

Pardiñas, A. F., P. Holmans, A. J. Pocklington, V. Escott-Price, S. Ripke, N. Carrera, S. E. Legge, S. Bishop, D. Cameron, M. L. Hamshere, et al., 2018 Common schizophrenia alleles are enriched in mutation-intolerant genes and in regions under strong background selection. Nature Genetics 50: 381–389.

Schumacher, F. R., A. A. Al Olama, S. I. Berndt, S. Benlloch, M. Ahmed, E. J. Saunders, S. Dadaev, D. Leongamornlert, E. Anokian, C. Cieza-Borrella, et al., 2018 Association analyses of more than 140,000 men identify 63 new prostate cancer susceptibility loci. Nature Genetics 50: 928–936.

Scott, R. A., L. J. Scott, R. Mägi, L. Marullo, K. J. Gaulton, M. Kaakinen, N. Pervjakova, T. H. Pers, A. D. Johnson, J. D. Eicher, et al., 2017 An expanded genome-wide association study of type 2 diabetes in Europeans. Diabetes 66: 2888–2902.

Martin, F. J., M. R. Amode, A. Aneja, O. Austine-Orimoloye, A. Azov, I. Barnes, A. Becker, R. Bennett, A. Berry, J. Bhai, et al., 2023 Ensembl 2023. Nucleic Acids Research 51: D933–D941.

Baumdicker, F., G. Bisschop, D. Goldstein, G. Gower, A. P. Ragsdale, G. Tsambos, S. Zhu, B. Eldon, E. C. Ellerman, J. G. Galloway, et al., 2022 Efficient ancestry and mutation simulation with msprime 1.0. Genetics 220: iyab229.

